# Inositol hexakisphosphate Functions as a Cofactor and Modulator of ADAR1 Activity

**DOI:** 10.64898/2026.01.17.699700

**Authors:** Francisco Venegas-Solis, Mengsi Lu, Roni Cohen-Fultheim, Felix Bender, Valeria Fedeli, Takuya Masunaga, Mikako Fujita, Thomas Zillinger, Sabine Schneider, Andreas Kaufmann, Adolfo Saiardi, Erez Y. Levanon, Henning Jessen, Stefan Bauer

**Affiliations:** Institute of Immunology, Philipps-Universität Marburg, Biomedizinisches Forschungszentrum Marburg, Marburg, Germany; Institute of Organic Chemistry & CIBSS - Centre for Integrative Biological Signalling Studies, University of Freiburg, Freiburg, Germany; Mina and Everard Goodman Faculty of Life Sciences, Bar-Ilan University, Ramat Gan, Israel; University of Bonn, University Hospital Bonn, Institute of Clinical Chemistry and Clinical Pharmacology, Bonn, Germany; Laboratory for Molecular Cell Biology, University College London, London, United Kingdom; Medicinal and Biological Chemistry Science Farm Joint Research Laboratory, Faculty of Life Sciences, Kumamoto University, 5-1 Oe-honmachi, Chuo-ku, Kumamoto, Kumamoto, Japan; Ludwig-Maximilians University Munich, Department of Chemistry, Munich, Germany

## Abstract

Adenosine-to-inosine (A-to-I) RNA editing by ADAR1 is a key post-transcriptional modification, and mutations in ADAR1 lead to Aicardi–Goutières syndrome (AGS), an autoimmune disorder. Despite its biological and clinical relevance, the regulation of ADAR1 activity remains incompletely understood. Using a combination of biochemical approaches, inositol-pentakisphosphate 2-kinase (IPPK)-knockout cells, molecular dynamics simulations, and a cell-permeable inositol hexakisphosphate (IP_6_) prodrug (Pro-IP_6_), we demonstrate that IP_6_ depletion drastically reduces global RNA editing, while supplementation with Pro-IP_6_ restores and even enhances editing levels. Furthermore, we identify the C6-phosphate of IP_6_ as a critical determinant of ADAR1 catalytic efficiency, functioning within a hydrogen-bonding network that indirectly coordinates a Zn²⁺-ion. Finally, we show that the AGS-associated ADAR1 mutation N907S impairs RNA editing activity, most likely by altering the hydrogen-bond interaction network linking IP_6_ to the ADAR1 catalytic center. Together, these findings identify IP_6_ as an essential cofactor and regulator of ADAR1 activity and highlight cofactor availability and interaction networks as strategies for therapeutically modulating RNA editing.

## Introduction

Adenosine-to-inosine (A-to-I) RNA editing is a crucial post-transcriptional modification mediated by adenosine deaminases acting on RNA (ADAR) proteins and influences RNA splicing, stability, and translation (*1*). The ADAR family comprises two isoforms of ADAR1 (interferon-inducible p150 and constitutively expressed p110), ADAR2, and ADAR3 (which is catalytically inactive). Most A-to-I editing events are mediated by ADAR1, primarily within repetitive elements such as *Alu* sequences. In addition to its role in gene expression regulation, RNA editing by ADAR1 is essential for distinguishing self from non-self RNA and for preventing aberrant type I interferon (IFN) production and cell death through innate immune dsRNA sensors such as melanoma differentiation-associated gene 5 (MDA5), protein kinase R, oligoadenylate synthases, and Z-DNA/RNA binding protein 1 (*2–7*).

Dysregulation of ADAR1 activity contributes to a variety of diseases. For example, mutations in *ADAR1* cause Aicardi–Goutières syndrome (AGS), a childhood autoimmune disorder characterized by chronic inflammation and an elevated type I IFN signature (*8*). In addition, mutations in ADAR1 have also been linked with bilateral striatal necrosis (BSN) (*9*) and dyschromatosis symmetrica hereditaria (DSH) (*10*). Beyond genetic mutations, altered RNA editing has also been associated with inflammatory and autoimmune diseases, including type 1 diabetes (*11*, *12*). Moreover, elevated ADAR1 expression or hyperediting is important in cancer progression (*13*). Depending on the virus, ADAR1 editing activity can also exert either proviral or antiviral effects (*14*).

In mice, *Adar* deficiency is embryonically lethal and causes a strong induction of IFNs (*15*, *16*). This phenotype is rescued by the depletion of immune dsRNA sensors (*2*, *6*, *7*, *17*, *18*). At the cellular level, conditional Adar1 deletion leads to the loss of several immune cell populations, including regulatory T cells (Foxp3-Cre), CD103⁺ dendritic cells (CD11c-Cre), alveolar macrophages (CD11c-Cre), and both immature and CD23⁺ mature recirculating B cells (Cd19-Cre) (*19–21*).

Despite its biological and clinical relevance, the regulation of ADAR proteins activity and expression/protein levels remains incompletely understood. The best-characterized mechanism involves the interferon-inducible ADAR1 p150 isoform (*22*), which connects immune activation to RNA editing. Additional regulatory layers include post-translational modifications such as SUMOylation and phosphorylations (*23*, *24*). Furthermore, acidic pH (*25*) and protein-protein interactions can modulate editing activity. One of these protein-protein interactions that stands out and modulates the editing activity is the formation of homodimers and heterodimers between ADAR1, ADAR2 and ADAR3 proteins (*26–28*).

Another regulatory layer of ADAR proteins is inositol hexakisphosphate (IP_6_). IP_6_ was described in 2005 as an essential cofactor for the correct folding and editing activity of human ADAR2 (*29*). Recently, cryo-EM structures of human ADAR1 revealed that IP_6_ is also a cofactor for this enzyme (*28*). In addition, mutations affecting IP_6_ synthesis in *Caenorhabditis elegans*, specifically in Inositol polyphosphate multikinase (*ipmk-1*) and inositol pentakisphosphate 2-kinase (*ippk-1*), have been reported to trigger antiviral responses. These mutations result in loss of ADAR-mediated RNA editing, leading to enhanced RNA interference, activation of the viral RNA sensor dicer-related helicase (DRH-1), and subsequent induction of the unfolded protein response (UPR) (*30*). However, a direct dependence of RNA editing by ADAR1 or ADAR2 on IP_6_ in mammalian cells has not yet been reported.

Here, we demonstrate that ADAR1 protein levels and RNA editing activity in mammalian cells depend on the cofactor IP_6_, and that changes in its metabolism lead to differential editing outcomes. Furthermore, using a combination of biochemical, cellular, and molecular dynamics approaches, we show that IP_6_ maintains a hydrogen-bond interaction network that indirectly coordinates Zn²⁺-ion positioning within the ADAR1 catalytic center, thereby controlling RNA editing efficiency. These findings indicate that IP_6_ is not only a structural cofactor required for ADAR1 folding, but also a dynamic regulator of its catalytic activity. Finally, we demonstrate that the AGS-associated ADAR1 mutation N907S markedly reduces RNA editing activity, most likely by disrupting the hydrogen-bond interaction network linking IP_6_ to the ADAR1 catalytic center, consistent with the loss-of-function phenotype observed in AGS.

## Results

### IPPK deficient cells have reduced levels of ADAR1 protein and RNA editing

IP_6_ is an essential cofactor of human ADAR2, required for proper folding and RNA editing activity (*29*). The residues of human ADAR2 that interact with IP_6_ have been identified through the crystal structure of the ADAR2-IP_6_ complex (PDB: 1ZY7). These residues are located within the deaminase domain, which is conserved across all ADAR family members (ADAR1-3) (Fig. S1A). Sequence alignment of the deaminase domains revealed that the IP_6_-interacting residues identified in ADAR2 are conserved in ADAR1 and ADAR3 (Fig. S1B), suggesting that, similar to ADAR2, ADAR1 also requires IP_6_ for correct folding and RNA editing activity. Furthermore, a cryo-EM structure of ADAR1 (PDB: 9B83) has recently confirmed the presence of IP_6_ (*28*), and interaction analysis verified most of the residues predicted by the sequence alignment (Fig. S1C and D).

Inositol-pentakisphosphate 2-kinase (IPPK) synthesizes IP_6_ by phosphorylating IP_5_ (specifically 1,3,4,5,6-IP_5_, the most abundant cellular isoform) (Fig. 1A and B; Fig. S2A) (*31*). In this manuscript, we use the term ‘IP_5_’ to refer specifically to the 1,3,4,5,6-IP_5_ isoform; other inositol pentakisphosphate isomers are named explicitly when used.

**Fig. 1.**
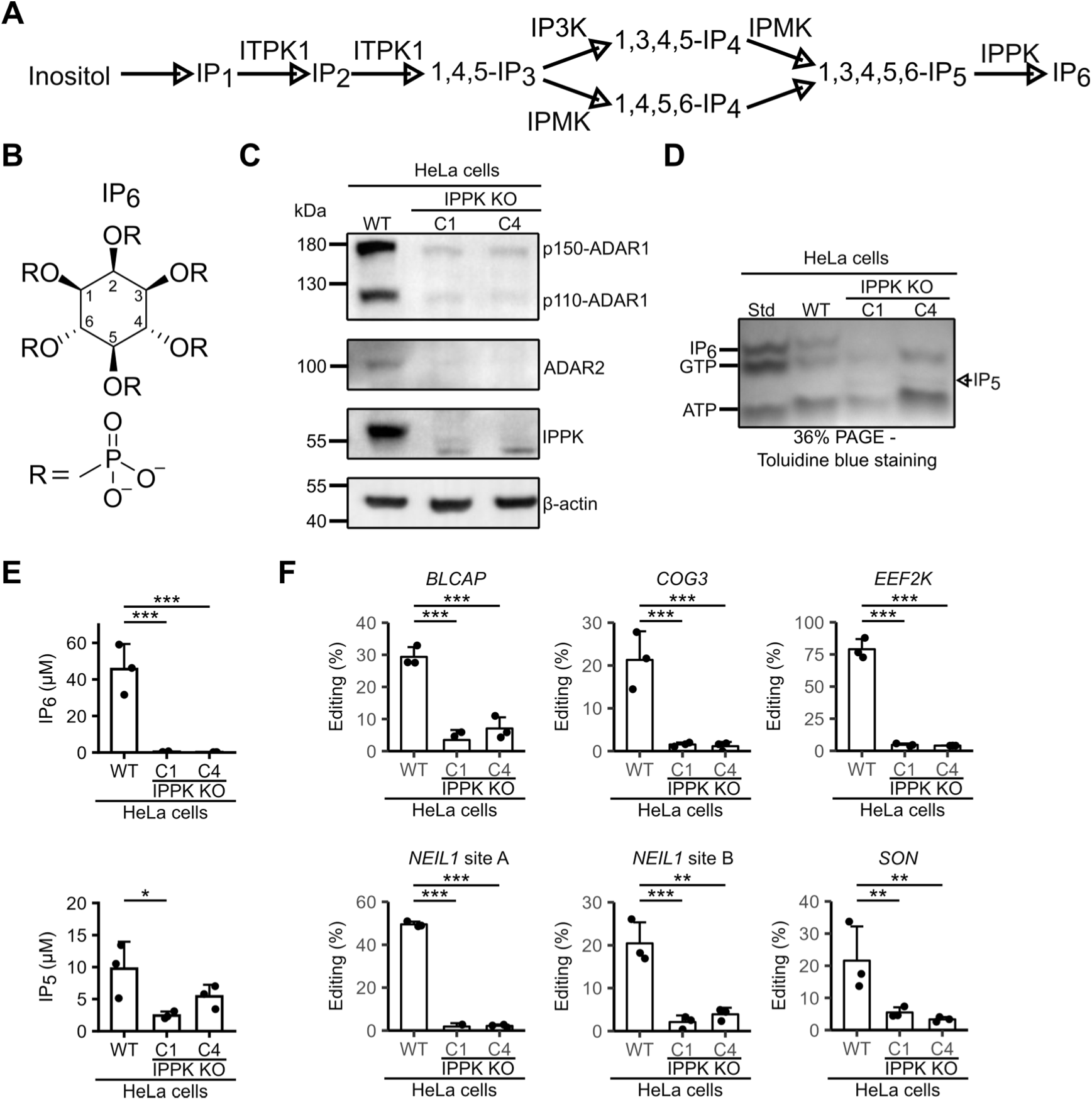
Dependency of ADAR1 and ADAR2 protein levels and activity on the cofactor inositol-hexaphosphate (IP_6_). (**A**) Schematic representation of the IP_6_ biosynthesis pathway mediated by inositol-tetrakisphosphate 1-kinase (ITPK1), Inositol polyphosphate multikinase (IPMK), Inositol-trisphosphate 3-kinase A (IP3K) and Inositol-pentakisphosphate 2-kinase (IPPK). (**B**) Structure of IP_6_. (**C**) Western blot analysis of ADAR1, ADAR2, IPPK and β-actin in HeLa cells and two HeLa IPPK KO clones (C1 and C4. (**D**) Detection of IP_6_ (GTP, ATP) levels in HeLa cells and HeLa IPPK KO (C1 and C4) cells utilizing 36% polyacrylamide gel electrophoresis followed by toluidine blue staining. (**E**) Quantification of IP_6_ and IP_5_ in HeLa WT and HeLa IPPK KO cells(C1 and C4) (n=3) using CE-MS. (**F**) RNA editing analysis of ADAR1 and ADAR2 editing sites in genes *BLCAP*, *COG3*, *EEF2K*, *NEIL1* and *SON* from HeLa WT cells or HeLa IPPK KO cells (C1 and C4) (n=3). Each data point represents one experiment. Statistical significance was analyzed using one-way ANOVA with Dunnett’s (E) post hoc test, *P < 0.05, **P < 0.01, or ***P < 0.001 compared to the WT cells.

To determine whether IP_6_ levels regulate A-to-I editing by ADAR1, we generated IPPK-deficient HeLa (HeLa IPPK KO) cells using CRISPR/Cas9. We found that IPPK KO cells lost ADAR2 protein and exhibit a dramatic reduction in ADAR1 protein level (p110 and p150 isoforms) (Fig. 1C) compared to WT cells. Furthermore, using polyacrylamide gel-toluidine staining and Capillary Electrophoresis-Mass Spectrometry (CE-MS) (*32–34*), we confirm that IPPK KO cells lost IP_6_, whereas IP_5_ was still present (Fig. 1D and 1E).

Regarding RNA editing, we analyzed six known editing sites from five genes (*BLCAP*, *COG3*, *EEF2K*, *NEIL1*, and *SON*; Table S1) (*35–38*) and observed that IPPK KO cells showed a drastic decrease in RNA editing in all the targets examined (Fig. 1F and Fig. S2B).

Overall, these results indicate that both ADAR1 protein and RNA editing activity are strongly dependent on IP_6_.

### Pro-IP_6_ treatment rescued ADAR1 protein level and activity in the IPPK KO cells

To determine the extent to which IP_6_ regulates ADAR1 protein levels and activity, we reintroduced IP_6_ into HeLa IPPK KO cells using a cell membrane-permeable prodrug IP_6_ (Pro-IP_6_) (*39*) (Fig. 2A). Pro-IP_6_ treatment (for 24 h) effectively restored intracellular IP_6_ levels and partially rescued ADAR1 and 2 proteins in IPPK KO cells (Fig. 2B and C). Of note, Pro-IP_6_ treatment (10 µM) produced an increase of IP_6_ levels and ADAR1/2 levels in WT cells (Fig. 2B and C; Fig. S3A).

**Fig. 2.**
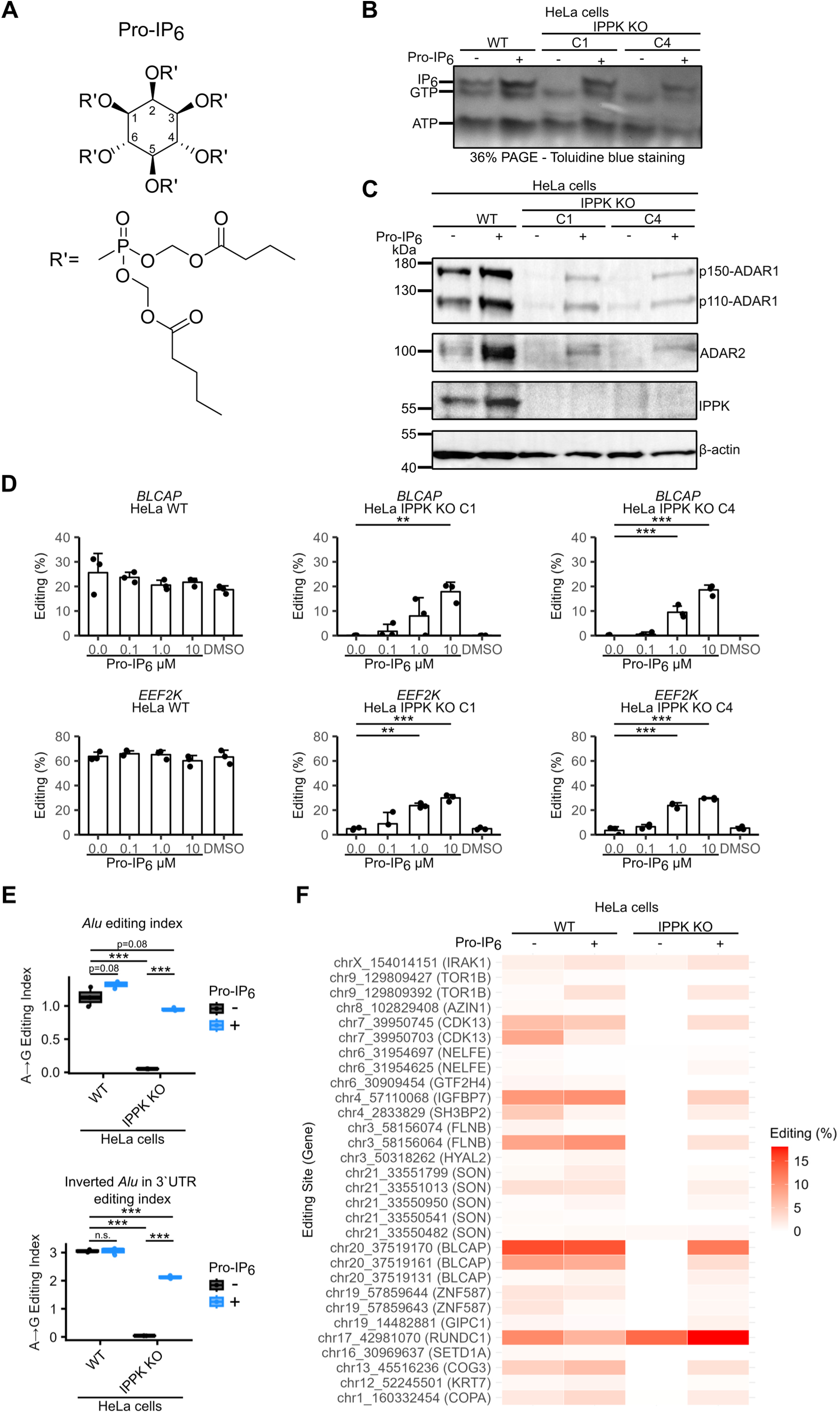
Pro-IP_6_ treatment restores RNA editing in HeLa IPPK KO cells. (**A**) Structure of butyryloxymethyl-modified IP_6_ (Pro-IP_6_). (**B**) IP_6_ levels in HeLa cells and HeLa IPPK KO cells C1 and C4 treated or untreated with 10 µM Pro-IP_6_ for 24 h were analyzed using 36% polyacrylamide gel electrophoresis and toluidine blue staining. (**C**) Western blot analysis of ADAR1, ADAR2, IPPK and β-actin in HeLa cells and HeLa IPPK KO C1 and C4 cells not treated or treated with 10 µM Pro-IP_6_ for 24 h. (**D**) Editing analysis of *BLCAP* and *EEF2K* editing sites in RNA from HeLa WT cells and HeLa IPPK KO C1 and C4 cells (n=3) untreated or treated with different concentrations of Pro-IP_6_ for 24 h. (**E**) *Alu* editing index and inverted *Alu* in 3’UTR editing index of HeLa cells (WT) and HeLa IPPK KO (C4) untreated or treated with10 µM Pro-IP_6_ for 24 h using NGS (n=3). (**F**) Editing analysis (heatmap) of some specific ADARs editing sites of HeLa cells (WT) and HeLa IPPK KO (C4) untreated or treated with10 µM Pro-IP_6_ for 24 h using NGS data. Each data point represents one experiment. Statistical significance was analyzed using one-way ANOVA with Dunnett’s (D) or Tukey’s (E) post hoc test, *P < 0.05, **P < 0.01, or ***P < 0.001 compared to the WT cells without treatment (D) or cells untreated with Pro-IP_6_ (E).

Importantly, Pro-IP_6_ treatment partially restored *BLCAP* and *EEF2K* editing levels in IPPK KO cells (Fig. 2D and Fig. S3B) in a dose-dependent manner, while WT cells showed no change in editing levels upon Pro-IP_6_ treatment.

To explore the global effects of IP_6_ depletion and Pro-IP_6_ treatment on RNA editing, we performed NGS on HeLa WT and HeLa IPPK KO cells, with or without Pro-IP_6_ treatment. Differential expression analysis identified a subset of significantly regulated genes (adjusted P < 0.05, |log₂FC| > 1), which were subjected to Gene Ontology (GO) Biological Process and Kyoto Encyclopedia of Genes and Genomes (KEGG) pathway enrichment analyses. Comparative gene expression profiling revealed marked differences between WT and IPPK KO cells. Among the biological processes enriched upon IP_6_ depletion were small GTPase–mediated signal transduction and muscle system processes. In parallel, KEGG pathway analysis identified enrichment of pathways including oxytocin signaling and human papillomavirus infection (Fig. S3C-E). These changes could result from the loss of ADAR-mediated RNA editing or from broader effects of IP_6_ depletion, since IP_6_ is a cofactor of additional proteins such as mTORC2 (*40*), CK2 (*41*), Ku70–Ku80 heterodimer (*42*), Pds5B (*43*), ICM (*44*) and GLE1 (*45*).

Since the loss of ADAR1 can trigger a type I interferon (IFN) signature (*17*), we analyzed whether IPPK KO cells display an upregulation of interferon-stimulated genes (ISGs) using NGS data. We did not observe a clear upregulation of ISGs in IPPK KO cells and in fact some ISGs were even downregulated compared to WT cells (Fig. S4). Based on these results, there is not a clear type I IFN signature in the IPPK KO cells.

To assess global RNA editing restoration by Pro-IP_6_ treatment in IPPK KO cells, we calculated the *Alu* editing index (*46*), a measure of the mean A-to-I editing across all *Alu* elements, and the inverted *Alu* 3′UTR editing index (*47*), defined as 3′UTR exons containing at least two *Alu* elements (each >200 bp) oriented on opposite strands. This metric captures editing activity within double-stranded RNA–forming 3′UTR structures, serving as a proxy for cytoplasmic A-to-I editing potential. Using these indices, we observed that Pro-IP_6_ treatment nearly completely restored global editing in IPPK KO cells and a partial rescue of cytoplasmic editing (Fig. 2E). Of note, Pro-IP_6_ treatment in WT cells caused a non-significant increase in editing, and a low level of editing persisted in KO cells (Table S2).

Analysis of A-to-I RNA editing across genomic regions revealed that intronic sequences contained the highest number of edited *Alu* elements, A→G mismatch sites, and total mismatches, consistent with the known enrichment of ADAR1 editing activity in intronic *Alu* repeats. IPPK KO cells exhibited decreased editing, which was restored by Pro-IP_6_ treatment. WT cells showed no significant change in editing levels upon treatment (Fig. S5A–C; Table S3).

For site-specific analysis, we quantified RNA editing levels at annotated CDS editing sites described by Gabay et al. (*48*) using REDItools 1.0 (*49*). Pro-IP_6_ treatment restored specific editing sites in IPPK KO cells and increased (e.g. TOR1B) or decreased (e.g. CDK13) editing at some sites in WT cells. Notably, three editing sites in SON, IRAK1 and RUNDC1 were also detected in untreated IPPK KO cells (Fig. 2F).

Regarding gene expression, PCA analysis showed no global rescue by Pro-IP_6_ treatment in IPPK KO cells (Fig. S6A). Nevertheless, we identified 51 genes whose expression shifted toward WT levels after treatment; the top 30 genes are visualized in Fig. S6B.

In general, these results demonstrate that loss of IP_6_ alters ADAR1/2 protein levels and RNA editing. Furthermore, Pro-IP_6_ treatment rescues ADAR1/2 protein and editing levels in IPPK KO cells and partially restores gene expression patterns. These findings indicate that IP_6_ availability modulates ADAR1 function. Despite the strong dependence on IP_6_, residual RNA editing in IPPK KO cells suggests that ADAR1 retains limited activity in the absence of IP_6_.

### A-to-I editing relationship with cellular IP_6_ levels

IP_6_ and IP_5_ levels vary depending on the cell type (Fig. 3A and Fig. S7A and B) (*50*). To determine whether RNA editing correlates with cellular inositol phosphates (IPs) levels, we quantified IP_6_ and IP_5_ in several cell lines from different origin (HeLa, HEK293, Caco-2, Jurkat, CAL-1 and THP-1) and in primary human peripheral blood mononuclear cells (hPBMCs) from 10 different donors. IPs levels were normalized to protein content and correlated to mean editing level of two different editing sites (*BLCAP* and *EEF2K*). For all cell lines (Fig. 3B and C) and hPBMCs (Fig. 3D-F), we observed a positive correlation between editing and IP_6_ levels, whereas no correlation was detected for IP_5_.

**Fig. 3.**
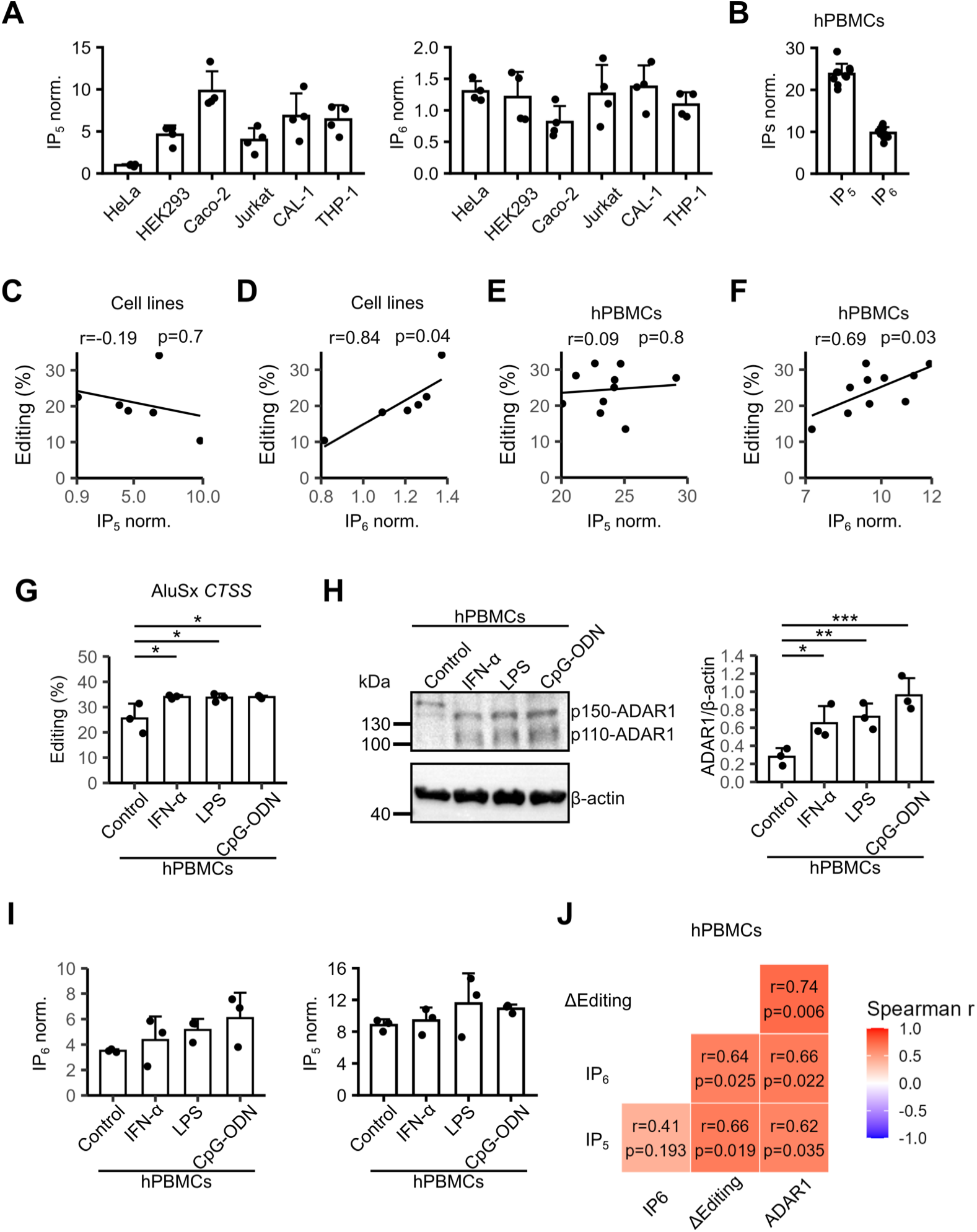
Cellular IP_5_ and IP_6_ levels and ADAR1-mediated RNA editing. (**A** and **D**) IP_6_ and IP_5_ (2-OH-IP_5_) levels normalized by protein amount in different cell lines (n=4) or hPBMCs were analyzed by CE-MS. Correlation of IP_5_ (2-OH-IP_5_) (**B** and **E**) and IP_6_ (**C** and **F**) levels normalized by protein amount against editing of *BLCAP* and *EEF2K* of six different cell lines (n=4) (**B** and **C**) and hPBMCs (n=10) (**E** and **F**). (**G**) Editing analysis of AluSx from *CTSS* gene in hPBMCs (n=3) untreated or treated with IFN-α (1 ng/mL), LPS (1 ng/mL) and CpG-ODN 2216 (1 µM) for 20 h. (23 editing sites with editing higher than 10% were considered). (**H**) Western blot analysis and quantification (n=3) of ADAR1 (p150+p110) normalized against β-actin in hPBMCs untreated or treated with IFN-α (1 ng/mL), LPS (1 ng/mL) and CpG-ODN 2216 (1 µM) for 20 h. Samples were analyzed for ADAR1 protein levels by western blotting. (**I**) Quantification of IP_6_ and IP_5_ in hPBMCs (n=3) treated or not with IFN-α (1 ng/mL), LPS (1 ng/mL) and CpG-ODN 2216 (1 µM) for 20 h, and IP_6_ or IP_5_ levels were quantified by CE-MS and normalized by protein amount. (**J**) Correlation analysis using the data from figures A-C. Each data point represents one donor or one experiment. For correlation analysis, Spearman (J) or Pearson (B, C, E and F) correlation coefficients were performed, and a P < 0.05 was considered significant. Statistical significance was analyzed using one-way ANOVA with Dunnett’s post hoc test, *P < 0.05, **P < 0.01, or ***P < 0.001 compared to the control cells.

To further investigate this relationship, we stimulated hPBMCs with type I IFN or type I IFN inducing stimuli (LPS, and CpG ODN 2216) to enhance RNA editing and evaluated whether this was accompanied by changes in IP_6_ and ADAR1 protein levels. A significant increase in editing (*Alu*Sx from *CTSS* (*51*, *52*), 23 editing sites analyzed) and ADAR1 protein was observed (Fig. 3G and 3H) upon stimulation, whereas compared to the control group a not significant increase of IP_6_ and IP_5_ was observed compared to the control group (Fig. 3I and Fig. S7C). Nevertheless, a strong significant positive correlation was observed between IP_6_ and ADAR1 protein levels and Δediting (Editing_Treatment_-Editing_Control_). Of note, IP_5_ levels also correlate positively with ADAR1 protein levels and Δediting (Fig. 3J).

Although total cellular IP_6_ and IP_5_ levels are generally considered stable (*53*), their actual availability has not been clearly defined, as these molecules can exist either protein-bound or free. Using size-exclusion filtration combined with IPs extraction, we determined that roughly half of the total IP_6_ and IP_5_ pool is protein-bound, while the other half remains “free” (completely free and/or lower affinity protein-IP_6_/_5_ complexes, where IPs were released during the extraction/purification process) (Fig. S7D). This indicates that fluctuations in IP_6_ and IP_5_ abundance may be more detectable when specifically assessing the free fraction rather than total levels.

As shown previously, Pro-IP_6_ treatment increased ADAR1 protein and global editing activity. To further clarify whether external modulation of IP_6_ levels directly affects editing, we treated HeLa WT and HeLa IPPK KO cells with Pro-IP_6_ and measured editing activity using an RNA editing-dependent dual-luciferase reporter assay (Fig. 4A). In this system, the Renilla luciferase coding sequence is followed by an RNA “bridge” containing a stop codon (UAG), with the A being a known editing site. When this site is edited by ADARs, translation proceeds to produce Firefly luciferase, making the Renilla/Firefly luciferase ratio a measure of global RNA editing. As expected, editing efficiency differed between WT and IPPK KO cells (Fig. 4B), and Pro-IP_6_ treatment increased RNA editing in both (Fig. 4C).

**Fig. 4.**
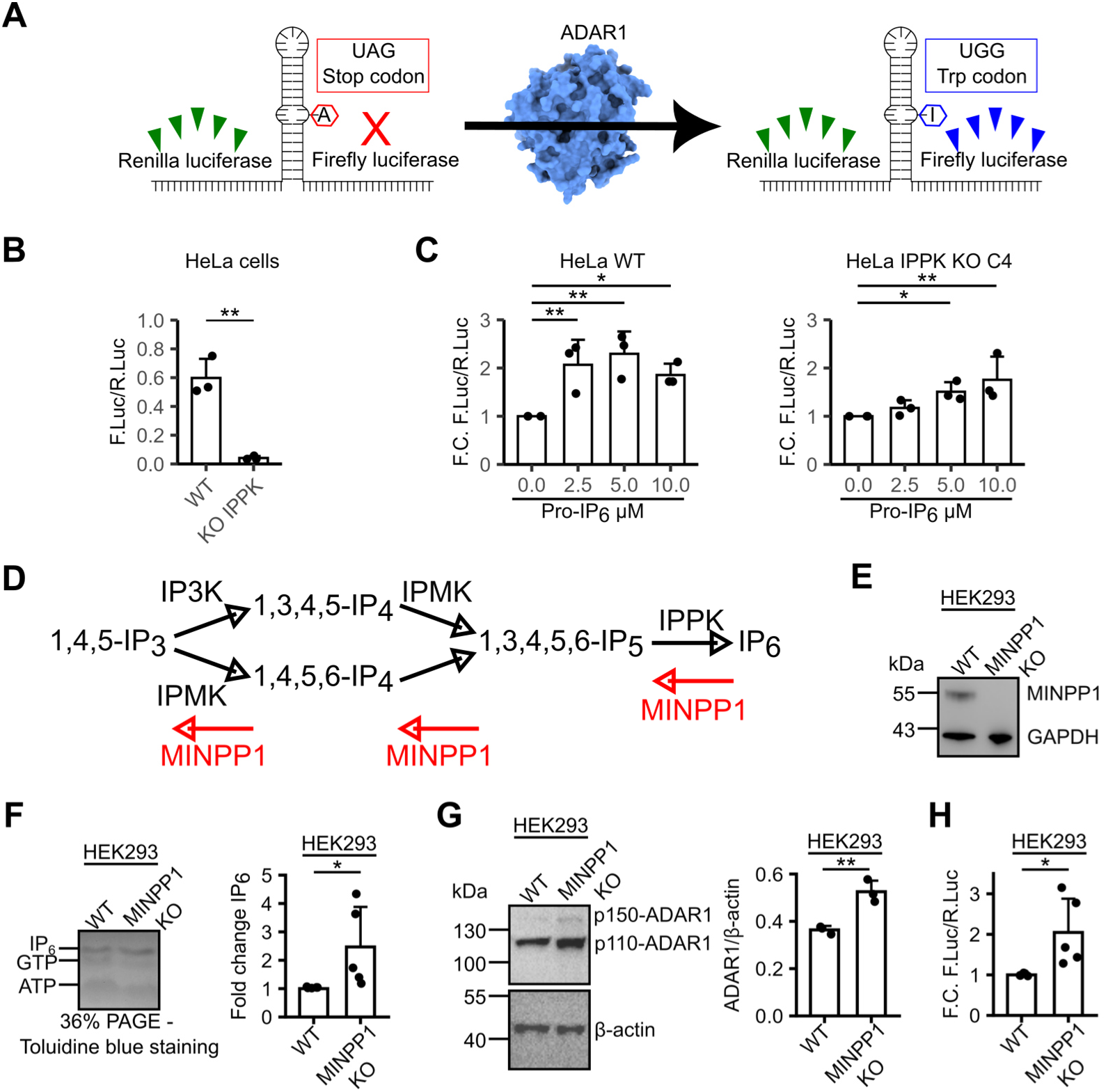
Modulation of RNA editing by IP_6_ levels. (**A**) Schematic representation of the RNA editing–dependent dual-luciferase reporter construct. (**B** and **C)** Editing analysis of HeLa cells and HeLa IPPK KO C4 (n=3) untreated or treated with different Pro-IP_6_ concentrations using the RNA editing–dependent dual-luciferase reporter assay. (**D**) Metabolic pathway of IP_6_ synthesis and degradation by IPMK, IPPK and multiple inositol polyphosphate phosphatase 1 (MINPP1). (**E**) Western blot analysis of MINPP and GAPDH in HEK293 cells and HEK293 MINPP KO cells. (**F**) IP_6_ analysis and quantification (n=5) in HEK293 cells and HEK293 KO MINPP cells by 36% polyacrylamide gel electrophoresis and toluidine blue staining. (**G**) Western blot analysis and quantification (n=3) of ADAR1 (p150+p110) and β-actin in HEK293 cells and HEK293 MINPP KO cells. (**H**) Editing analysis of HEK293 cells and HEK293 MINPP KO cells (n=3) using the RNA editing–dependent dual-luciferase reporter assay. Each data point represents one experiment. Statistical significance was analyzed using Student’s t test (H, L, M and N) or one-way ANOVA with Dunnett’s (I) post hoc test compared to WT cells without treatment, *P < 0.05, **P < 0.01, or ***P < 0.001.

Multiple inositol polyphosphate phosphatase 1 (MINPP1) hydrolyzes IP_6_, IP_5_ and IP_4_ to less phosphorylated inositol phosphates (Fig. 4D) (*54*). Cells deficient in MINPP1 exhibit increased IP_6_ levels compared to WT cells (*55*). To determine whether elevated IP_6_ enhances ADAR1 protein levels and RNA editing, we used HEK293 WT and MINPP1 KO cells (Fig. 4E) and quantified ADAR1 protein and editing activity using the dual-luciferase reporter assay. Consistently, MINPP1-deficient cells have higher IP_6_ and ADAR1 protein levels, accompanied by increased RNA editing (Fig. 4F–H).

Together, these results demonstrate that RNA editing and ADAR1 expression correlate strongly with intracellular IP_6_ levels. Moreover, external modulation of cellular IP_6_ abundance by Pro-IP_6_ enhances ADAR1 protein and RNA editing activity.

### IP_6_ and IP_5_ as cofactors of ADAR1

As mentioned before, we observed that ADAR1 abundance and activity was not completely lost in IPPK KO cells resulting in a residual RNA editing activity (Fig. 1). Based on these results, we hypothesized that another inositol phosphate might partially substitute for IP_6_ as a natural cofactor of ADARs. To test this hypothesis, we performed *in vitro* translation of ADAR1 in the presence of either IP_6_ or IP_5_ or no IPs (Fig. 5A).

**Fig. 5.**
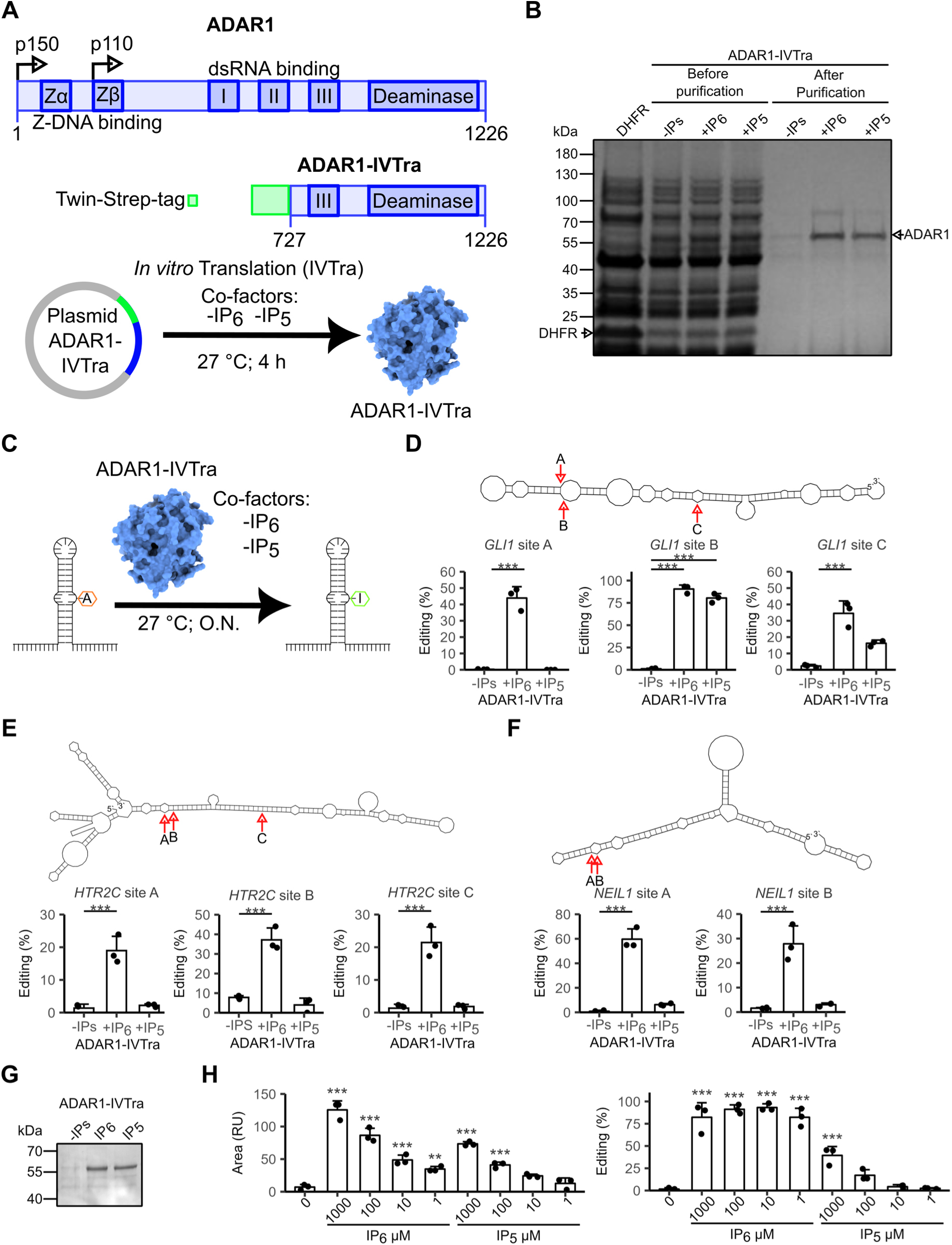
IP_6_ and IP_5_ as a cofactors for *in vitro* ADAR1. (**A**) Scheme of ADAR1 protein, *ADAR1 in vitro* translated construct (ADAR1-IVTra) and the experimental setup for protein generation. (**B**) *In vitro* synthesized ADAR1-IVtra using no IPs or IP_6_ or IP_5_ (2-OH-IP_5_), were analyzed by SDS-PAGE and Coomassie staining before and after purification with MagStrep “type3” XT beads. Dihydrofolate reductase (DHFR) was used as control of the *in vitro* translation synthesis. (**C**) Schematic representation of the *in vitro* editing reaction using ADAR1-IVTra and different RNAs (**D-F**) *In vitro* editing reaction (n=3) of *in vitro* transcribed *GLI1*, *HTR2C* and *NEIL1* dsRNA using ADAR1-IVTra; secondary structure of the dsRNAs fragments (editing sites are depicted by arrows and A, B and C) were predicted using RNAfold; editing levels were analyzed by cDNA synthesis follow up by PCR amplification and sanger sequencing. (**G**) ADAR1-IVTra proteins used in the *in vitro* reactions of (**D**-**F**) were analyzed by SDS-PAGE and Coomassie staining. (**H**) Quantification of ADAR1-IVTra protein using different concentrations of IP_6_ and IP_5_ (left graphic), and corresponding editing of *GLI1* B editing site after 1 h of *in vitro* editing reaction (right graphic). Each data point represents one experiment or one donor. Statistical significance was analyzed using one-way ANOVA with Dunnett’s post hoc test, *P < 0.05, **P < 0.01, or ***P < 0.001 compared to ADAR1-IVTra-no IPs.

We observed *in vitro* translated ADAR1 (ADAR1-IVTra) in all conditions tested but only ADAR1-IVTra with IP_6_ or IP_5_ was recovered after the purification using MagStrep “type3” XT beads (Fig. 5B). This suggests that inositol phosphates such as IP_5_ or IP_6_ facilitate proper folding/stabilization of ADAR1, enabling successful purification. We next evaluated the RNA editing activity of ADAR1-IVTra using synthetic dsRNAs corresponding to known ADAR targets (*GLI1*, *HTR2C*, and *NEIL1*) (*15*, *56–58*) (Fig. 5C, Fig. S8 and Table S1). The *in vitro* editing assays showed that ADAR1-IVTra-IP_6_ efficiently edited all sites tested, whereas ADAR1-IVTra-IP_5_ edited only 2 of 8 sites and with reduced efficiency (Fig. 5D–G and Fig. S9A).

For comparison, we also tested *in vitro* translated ADAR2, which showed a different pattern compared with ADAR1 production/recovery. While IP_6_ supported production/recovery and activity, IP_5_ did so to a lesser extent (Fig. S9B). *In vitro* editing assays demonstrated that ADAR2-IVTra-IP_6_ edited 3 of 8 sites, whereas ADAR2-IVTra-IP_5_ showed no detectable editing activity (Fig. S9D–F).

To further study whether IP_5_ can function as a cofactor of ADAR1, we synthesized ADAR1 *in vitro* with increasing concentrations of IP_5_ or IP_6_ and quantified both protein yield and editing activity on *GLI1* RNA at site B. Protein production was concentration-dependent for both cofactors, but IP_6_ supported ADAR1 production more efficiently. Regarding editing activity, all IP_6_ concentrations (1 mM to 1 µM) supported robust editing, whereas only the two highest IP_5_ concentrations (1 mM and 100 µM) resulted in detectable editing (Fig. 5H).

It is known that ADAR1-mediated RNA editing preferentially occurs at A:C mismatches rather than A:U pairs (*59*). From an energetic and structural perspective, we observed that editing efficiency increased when adenosine deamination resulted in a more stable RNA structure (Fig. S9H). This effect was typically associated with the closure of RNA loops in the secondary structure, suggesting that these adenosines are more accessible than those involved in perfect base pairing. Based on this correlation, we propose that ADAR1-IP_5_ editing predominantly occurs in RNAs containing loops that expose the adenosine to deamination and confer a favorable energetic change upon loop closure.

Overall, these results indicate that IP_5_ can partially substitute for IP_6_ as a cofactor under IP_6_-depleted conditions, supporting limited RNA editing activity.

### Analysis of the ADAR1-IP_6_ binding site

To examine the IP_6_-binding pocket of ADAR1 in more detail and to evaluate differences between IP_5_ and IP_6_ binding, we performed molecular dynamics (MD) simulations of the ADAR1 deaminase domain in complex with either IP_6_ or IP_5_. The stability of the complexes was assessed using root mean square deviation (RMSD), and the flexibility of individual residues was evaluated using root mean square fluctuation (RMSF). RMSD analysis indicated that the ADAR1-IP_5_ complex maintained a similar conformation compared with the ADAR1-IP_6_ complex (Fig. S10A). Consistently, RMSF analysis (n=3) indicated that the ADAR1-IP_5_ complex maintained a residue-level flexibility profile similar to that of ADAR1-IP_6_, with only minor differences at a few peak regions (Fig. S10B). Together, these results indicate that IP_5_ binding does not substantially alter the global structure or local dynamics of ADAR1.

With respect to direct interaction between ADAR1 and IP_6_ or IP_5_, we identified 15 residues forming 23 interactions (distance: 1.6 – 2.5 Å) with IP_6_ and 13 residues forming 19 interactions with IP_5_ (Fig. 6A). The interactions lost in the ADAR1-IP_5_ complex due to the absence of the C2-phosphate involved residues R917, K1158, Y1182 and K1186. MM-GBSA calculations (n=3, 50 ns per simulation) estimated the relative binding free energies of the two complexes and showed that ADAR1–IP_6_ has a substantially more favourable binding energy than ADAR1-IP_5_, suggesting a stronger association of IP_6_ under the simulated conditions (Fig. 6B).

**Fig. 6.**
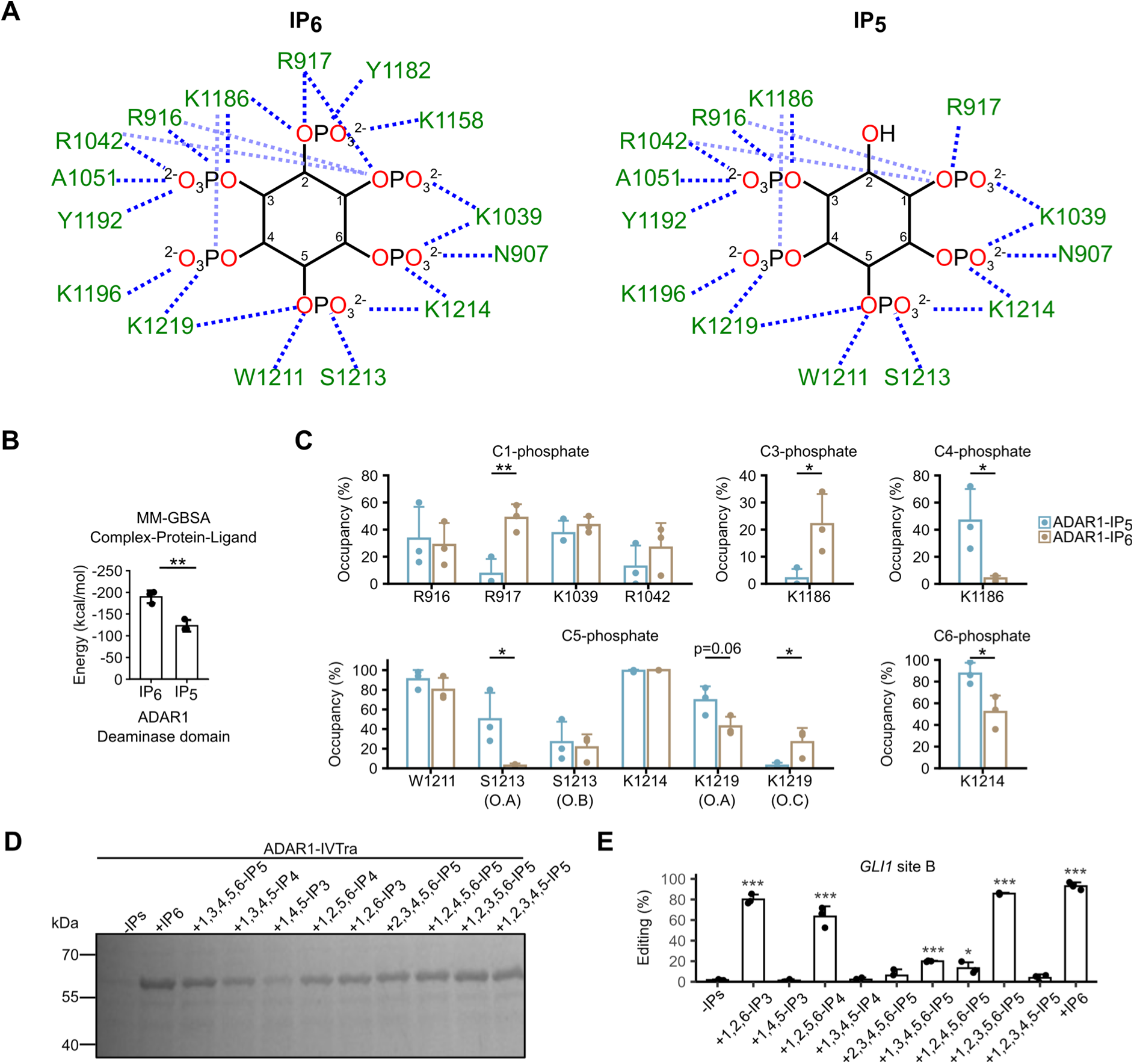
IP_6_ and IP_5_ binding analysis in ADAR1. (**A**) Interaction analysis based on molecular dynamics (MD) simulations between the editase domain of ADAR1 and IP_6_ or IP_5_. Interacting residues are shown in green, and interactions are represented by blue dashed lines. (**B**) MM-GBSA binding free energy calculations (n = 3) of ADAR1-IP_6_ and ADAR1-IP_5_ complexes based on MD simulations. (**C**) Hydrogen bond occupancy (n = 3) of specific interactions between ADAR1 and IP_5_ or IP_6_. (**D**) ADAR1-IVTra proteins synthesized in the presence of different inositol phosphates (IPs) were analyzed by SDS-PAGE followed by Coomassie staining. (**E**) *In vitro* editing reactions (n = 3) of the GLI1 dsRNA site B using ADAR1-IVTra with different IPs as cofactors. Each data point represents one individual experiment. Statistical significance was analyzed using Student’s t test (B and C) or one-way ANOVA with Dunnett’s post hoc test (E) compared to ADAR1-IVTra-no IPs. *P < 0.05, **P < 0.01, or ***P < 0.001.

We hypothesized that the lost of the C2-phosphate in IP_5_ could alter the dynamic of interactions involving the remaining phosphates. To evaluate this hypothesis, we analyzed the occupancy of specific interactions (defined as the percentage of simulation time during which an interaction was present) that were likely to be affected by the absence of the C2-phosphate (Fig. 6C). R917 interacts with the C1/C2-phosphates in the ADAR1-IP_6_ complex; in the ADAR1–IP_5_ complex, R917-C1-phosphate interaction was markedly reduced. K1186 interacts with the C2/C3/C4-phosphates of IP_6_; in the IP_5_ complex, this residue showed reduced occupancy with the C3-phosphate but increased occupancy with the C4-phosphate.

We also observed a slight shift of IP_5_ toward interactions involving the C5-phosphate compared with IP_6_. In this context, residues S1213 and K1219 displayed altered interaction patterns in the IP_5_ scenario. For interaction analysis, the three non-bridging oxygens of the C5-phosphate were designated O.A, O.B, and O.C. S1213 interacted with both O.A and O.B in the ADAR1-IP_5_ complex, whereas in the ADAR1-IP_6_ complex it interacted only with O.B. Similarly, K1219 interacted with O.A but not O.B in the ADAR1-IP_5_ complex, whereas in the ADAR1-IP_6_ complex it interacted with both oxygens. Given these changes and the spatial proximity of nearby residues, we also analyzed K1214, which interacts with both the C5- and C6-phosphates. K1214 showed more efficient interaction with the C6-phosphate when IP_5_ was bound (Fig. 6C). Alignment of the IP_6_ binding site of the two complexes revealed differences in interaction distances betweenC5/C6-phosphates of IP_6_ or IP_5_ and residues S1213 and K1214 (Fig. S10C).

The reduced catalytic activity of the ADAR1-IP_5_ complex, compared with the ADAR1-IP_6_ complex, is not only due to the differences in overall protein stability, but also to perturbation of the interaction network linking IP_6_ to the catalytic Zn²⁺-ion, caused by the absence of the C2 phosphate. This network comprises interactions between IP_6_, residues K1039, D908, and K1003, and ADAR1 catalytic center, including the Zn²⁺-ion coordinated by H910, C966, and C1036. Specifically, K1039, D908, and K1003 form an interaction chain that functions as a molecular “bridge” between IP_6_ and the Zn²⁺-ion. K1039 directly interacts with the C1/C6 phosphates of IP_6_, while K1003 and D908 interact with C1036 and H910, respectively, which in turn coordinate the Zn²⁺ ion (Fig. S10D,E).

To gain further insight into which phosphates from IP_6_ may affect ADAR1 stability and RNA editing activity, we performed *in vitro* translation in the presence of different inositol phosphates and assessed protein recovery and editing activity. With respect to protein production/recovery, we observed that all IPs tested were able to support ADAR1 protein generation (Fig. 6F). Interestingly, the most common isomers of IP_3_ (1,4,5-IP_3_) and IP_4_ (1,3,4,5-IP_4_) were those that presented a lower amount of purified protein compared to the other IPs.

Regarding RNA editing activity, editing at the A site of *GLI1* was detected only for ADAR1-IVTra-IP_6_ (Fig. S11A). At site B, five additional isomers showed editing activity (1,2,6-IP_3_; 1,2,5,6-IP_4_; 1,3,4,5,6-IP_5_; 1,2,4,5,6-IP_5_; 1,2,3,5,6-IP_5_) besides IP_6_ (Fig. 6G), while at site C, only the two isoforms 1,2,6-IP_3_ and 1,2,3,5,6-IP_5_ supported editing (Fig.S11A). Based on IP_5_ isomer results, loss of the C1/6-phosphates completely abolished editing activity, whereas loss of the C2/3-phosphates caused a considerable reduction compared to other isoforms and to IP_6_, which exhibited the highest editing efficiency at the *GLI1* B site. Interestingly, 1,2,6-IP_3_ and 1,2,5,6-IP_4_ isomers but not 1,4,5-IP_3_ and 1,3,4,5-IP_4_ showed editing comparable to IP_6_ at the B site. These findings suggest that C1/2/6-phosphates of IP_6_ are the most critical for maintaining a minimal RNA editing activity by ADAR1, with position C6 likely being the most important, due that C6-phosphate interacted directly with “bridge” that join the IP_6_ and the Zn^2+^-ion of the catalytic site.

We next hypothesized that exchange between IP_6_ and IP_5_ could be feasible, ADAR1-IVTra was generated with IP_6_ as cofactor, purified and used for *in vitro* editing in the presence of different IP_5_ concentrations, no IPs or IP_6_. For the controls (no IPs and IP_6_), no differences in *GLI1* editing were observed, but increasing concentrations of IP_5_ led to a reduction in *GLI1* editing (Fig. S11B and S11C) supporting the notion that IP_5_ can partially substitute IP_6_ in an exchange reaction.

In summary, these results demonstrate that specific phosphates of IP_6_ are essential for maintaining ADAR1 RNA editing activity, likely on one side by impacting on the correct folding and overall complex stability, as well as maintaining the required active site geometry through modulation of the hydrogen-bond interaction network linking IP_6_ to the ADAR1 catalytic center. Furthermore, our data indicate that exchange between IP_6_ and IP_5_ is biochemically feasible, albeit functionally suboptimal.

### ADAR1-N907S, an Aicardi-Goutières syndrome mutation

In the previous section, we identified the C6-phosphate of IP_6_ as one of the key determinants of ADAR1 RNA editing activity, playing a central role within a network connecting IP_6_ to the catalytic Zn²⁺-ion. According to our analysis of the IP_6_ binding site, this phosphate interacts with three residues in ADAR1: N907, K1039, and K1214 (Fig. 7A). Notably, a mutation at N907 (N907S) has been associated with AGS (*60*). To investigate the effect of this mutation, we introduced it into the ADAR1-IVTra construct and assessed both, protein recovery and RNA editing activity. For comparison, we also included the inactive E912A mutant (*8*), the AGS-associated G1007R mutant (*8*), and the hyperediting E1008A mutant (*28*) (Fig. 7B).

**Fig. 7.**
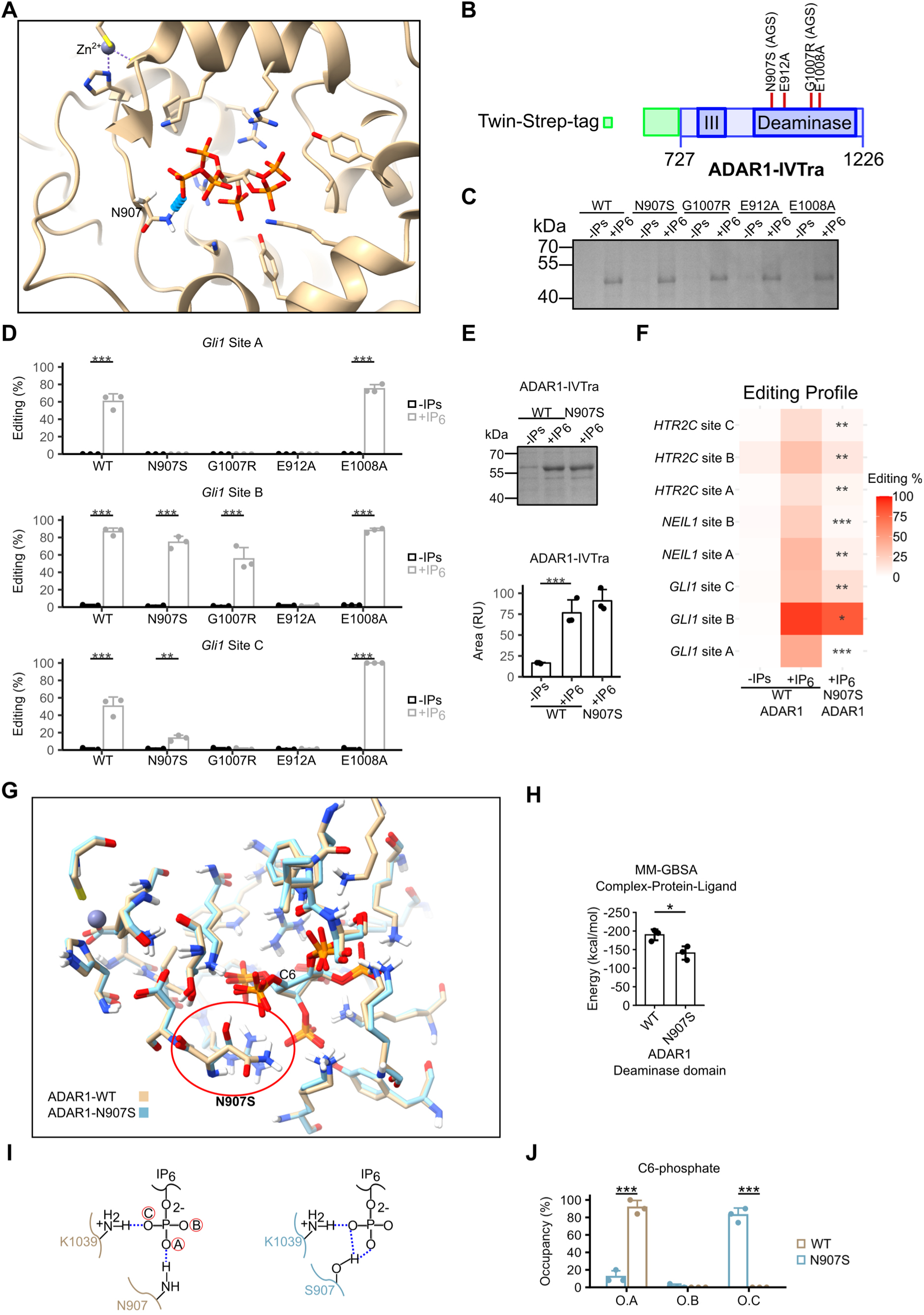
ADAR1-N907S editing and mechanism. (**A**) Structural overview of the localization of the residue N907 in ADAR1 (PDB: 9B83). (**B**) Schematic representation of ADAR1-IVTra construct and the localization of two mutations associated with AGS (N907S and G1007R), one catalytic inactive mutation (E912A) and one hyper-editing mutation (E1008A). (**C**) ADAR1-IVtra WT and mutant proteins were synthesized with no IPs or IP_6_ and visualized by SDS-PAGE and Coomassie staining. (**D**) *In vitro* editing levels (n=3) of *GLI1* dsRNA using ADAR1-IVTra WT and mutants synthesized with no IPs or IP_6_. (**E**) ADAR1-IVtra WT (no IPs or IP_6_) and N907S-IP_6_ were analyzed and quantified by SDS-PAGE and Coomassie staining. (**F**) Editing profile (heatmap) of ADAR1-IVTra WT (no IPs and IP_6_) and ADAR1-IVTra N907 (IP_6_) for 8 editing sites in *GLI1, NEIL1* and *HTR2C* dsRNA. (**G**) Close-up view of the IP_6_ binding site from the structural alignment of ADAR1-WT and ADAR1-N907S. Both structures were generated by MD simulations. The red circle marks the N907S mutation. (**H**) MM-GBSA binding free energy calculations (n = 3) of ADAR1-WT and ADAR1-N907S based on MD simulations. (**I**) Schematic representation of hydrogen bonds formed between N907 or S907 and the C6-phosphate of IP_6_ in ADAR1-WT or ADAR1-N907S, respectively. (**J**) Hydrogen bond occupancy (n = 3) of N907 or S907 with the C6-phosphate in ADAR1-WT or ADAR1-N907S. Each data point represents one experiment. Statistical significance was analyzed using Student’s t test (H and J) or one-way ANOVA with Dunnett’s (E and F) or Tukey’s (D) post hoc test, *P < 0.05, **P < 0.01, or ***P < 0.001 compared to ADAR1-IVTra-no IPs.

With respect to protein generation by *in vitro* translation, none of the mutations appeared to affect ADAR1 recovery compared to ADAR1-IVTra WT (Fig. 7C). Regarding *GLI1* editing, both ADAR1-IVTra WT and the E1008A mutant showed editing at all sites, with the E1008A mutant displaying higher editing efficiency at site C. The E912A mutant had no editing activity at any site tested, while the AGS-associated mutants (N907S and G1007R) exhibited no editing at site A, reduced editing at site B, and in the case of N907S, residual editing at site C, although at lower levels than the WT (Fig. 7D).

To gain a broader understanding of the impact of the N907S mutation on RNA editing, we performed *in vitro* editing reactions using *NEIL1* and *HTR2C* dsRNA and compared the editing activity between the N907S mutant and the WT protein. These assays confirmed that the mutant is produced and recovered at the same level than WT ADAR1-IVTra but showed markedly reduced editing activity (Fig. 7E and F; Fig. S12A-C).

To investigate structural consequences of the N907S substitution, we performed MD simulations of ADAR1-WT and ADAR1-N907S. RMSD analysis (n=3) indicated that the ADAR1-N907S adopts a slightly more stable and less fluctuating global conformation compared with ADAR1-WT (Fig. S12D). RMSF analysis (n=3) showed largely similar residue-level flexibility profiles between the two proteins, with only minor differences at specific regions (Fig. S12E). These results suggest that the N907S mutation does not induce global structural destabilization or major changes in local dynamics and may instead confer modest overall stabilization.

The most pronounced structural difference between ADAR1-WT and ADAR1-N907S was observed in the positioning of the side chain at residue 907 (Fig. 8A). MM-GBSA calculations (n = 3, 50 ns per simulation) revealed that ADAR1-WT binds IP_6_ with a substantially more favourable free energy compared with ADAR1-N907S, indicating reduced IP_6_ affinity upon mutation (Fig. 8B).

To analyze interactions at the mutation site in more detail, the three non-bridging oxygens of the C6 phosphate were designated O.A, O.B, and O.C. In ADAR1-WT, N907 primarily interacted with O.A, whereas in ADAR1-N907S, S907 interacted predominantly with O.C (Fig. 8C and D). Moreover, S907 displayed an oscillatory motion between O.A and O.C, introducing increased local dynamics relative to the more stable N907 interaction. This altered behavior was accompanied by an elongation of the interaction between K1039 and the C6 phosphate (O.C) (Fig. S12F). This mutation appears to increase the local dynamics and impacts the interactional network linking IP_6_ to the ADAR1 catalytic center.

Overall, these data demonstrate that the AGS-associated N907S mutation impairs ADAR1 RNA editing activity, likely by increasing the dynamics of the hydrogen-bond interaction network linking IP_6_ to the ADAR1 catalytic center, rather than by globally destabilizing the protein. This mechanistic insight provides a structural and functional explanation consistent with the pathogenic role of the N907S mutation in AGS.

## Discussion

IPs are a family of intracellular messengers that play key roles in diverse cellular processes, including cell growth, energy metabolism, vesicular trafficking, and immune signaling (*50*). One of this cellular process is A-to-I RNA editing, in which IP_6_ is an essential cofactor for ADAR1 and ADAR2, contributing to protein folding and RNA editing activity (*28*, *29*). However, the extent to which ADAR1/2 proteins depend on IP_6_ in mammalian cells has not been previously investigated.

Two main pathways contribute to IP_6_ synthesis. The lipid-derived pathway generates IP_3_ from phosphatidylinositol (3,4,5)-trisphosphate, while the cytosolic pathway produces IP_3_ via the activity of inositol-tetrakisphosphate 1-kinase (ITPK1). In both cases, further phosphorylation steps catalyzed by (inositol polyphosphate multikinase) IPMK and Inositol-pentakisphosphate 2-kinase (IPPK) lead to the formation of IP_6_ (*61*). Here, we show that IPPK-deficient cells almost completely lose their capacity for ADAR-mediated RNA editing, primarily as a consequence of a drastic reduction in ADAR1 levels. A similar observation was recently reported in HEK293T cells, where IPPK deficiency resulted in reduced levels of both ADAR1 and ADAT1. In that study, partial loss of these proteins was attributed to proteasome-mediated degradation, and impaired tRNA editing by ADAT1 was also observed (*30*).

The residual editing activity in these cells could be explained by IP_5_ partially substituting for IP_6_ as a cofactor for ADAR1. Alternatively, there might exist a secondary route for IP_6_ synthesis; however, our results suggest that it produces only low levels of IP_6_, with IP_5_ remaining more abundant than residual IP_6_ in IPPK-deficient cells.

The abundance of individual IP species varies across cell types. For example, some human cells contain higher levels of IP_5_ than others, whereas IP_6_ is generally the most abundant IP species (*32*, *50*). In this study, we found that certain cancer cell types and hPBMCs exhibit higher IP_5_ than IP_6_ levels. This raises intriguing questions about whether an ADAR1–IP_5_ complex might form physiologically under specific cellular conditions.

Treatment with Pro-IP_6_ partially restored RNA editing in IPPK-deficient cells, although ADAR1 protein was only modestly recovered. This limited effect could be due to the short treatment duration (24 h); prolonged exposure might lead to stronger restoration, approaching wild-type levels. It is worth noting that the observed protein levels changes and the recovery by Pro-IP_6_ treatment are not necessarily solely due to RNA editing by ADAR1, since IP_6_ is also a cofactor of other proteins and a signaling molecule (*40–45*).

IP_6_ is known to regulate Bruton’s tyrosine kinase (Btk) in B cell receptor (BCR) signaling and is also required for the proper activation of regulatory T (Treg) cells, including their differentiation into RORγt⁺ and tissue-resident subsets (*62*, *63*). Thus, loss of IP_6_ could lead to reduced RNA editing and impaired ADAR1 function, potentially explaining defects in immune homeostasis. This hypothesis is supported by a study showing that Foxp3-specific depletion of *Adar* in mice results in reduced numbers of Treg cells (*21*). Together, these findings indicate that IP_6_ levels, which influence ADAR1 protein levels and activity, may play an important role in cell differentiation and immune regulation.

Loss of ADAR activity results in reduced RNA editing, particularly within *Alu* elements, which can in turn activate MDA5 and drive type I interferon (IFN) production (*2*, *6*, *18*). A recent study reported a type I IFN signature in IP_6_-deficient cells based on the expression of a limited set of IFN-related genes (*30*). In contrast, analysis of our NGS data did not reveal a clear type I IFN response upon IP_6_ depletion, despite a pronounced reduction in ADAR1 protein. These differences may reflect context-dependent effects of inositol phosphate metabolism on innate immune signaling. Notably, the kinases IPPK, PPIP5K1, and PPIP5K2 (which convert IP_5_ to IP_6_ and IP_7_) have been shown to be critical for RIG-I-mediated interferon induction in HEK293 cells (*64*). Together, these observations suggest that the relationship between IP_6_ depletion and type I interferon activation is complex and likely influenced by cell type, experimental context, and the specific innate immune pathway engaged.

Regarding the dynamics of IP_6_ and IP_5_ production, these metabolites are considered relatively stable, with changes occurring only over long timescales compared to less phosphorylated IP species, typically under conditions such as phosphate starvation (*50*, *65*). Nevertheless, it is important to distinguish between total and free (unbound) IP_6_ pools, as changes in the unbound fraction could occur more dynamically and be functionally relevant. In our study, we found that IP_6_ levels correlate with editing activity in both tumor cell lines and primary cells.

Pontocerebellar hypoplasia (PCH) is a neurodevelopmental disorder caused by loss-of-function mutations in the *MINPP1* gene, which encodes multiple inositol-polyphosphate phosphatase 1 (MINPP1) (*55*, *66*). MINPP1 is responsible for dephosphorylating IP_6_, and its deficiency leads to intracellular accumulation of IP_6_ (*55*). In agreement with this, we found that MINPP1-deficient cells not only exhibit elevated IP_6_ levels but also show increased ADAR1 protein and activity. These results raise the possibility that patients with PCH may display enhanced RNA editing due to increased ADAR1 protein, potentially contributing to disease mechanisms.

We confirm that IP_6_ supports the proper folding of both ADAR1 and ADAR2, as previously reported (*28*, *29*). Interestingly, other IPs can also assist ADAR1 folding, but although they support only minimal RNA editing activity. Indicating that correct folding is necessary but not sufficient for full catalytic function. Despite IP_6_ being “buried” within the ADAR core, our *in vitro* assays show that exchange between IP_6_ and IP_5_ is feasible, suggesting that the ADAR1 complex may exist in a more dynamic cofactor-binding environment than previously appreciated.

Based on the crystal structure of the ADAR2–IP_6_ complex, it was suggested that the C6-phosphate interacts indirectly with Zn²⁺-ion through a hydrogen-bond network involving K519, D392, and K483, thereby contributing to proper positioning of Zn²⁺-ion within the catalytic center (*29*). In this study, we show that, like ADAR2, ADAR1 possesses a hydrogen-bond network in which the C1 and C6 phosphates of IP_6_, together with residues K1039, K1003, and D908, play central roles. This network appears to indirectly impact on RNA editing activity by stabilizing Zn²⁺-ion positioning within the catalytic center.

The reduced RNA editing activity observed in ADAR1 in the presence of alternative inositol phosphates, despite preserved protein folding, likely reflects the ability of these IPs to meet minimal structural requirements without maintaining the integrity or rigidity of the hydrogen-bond interaction network. Therefore, proper folding of the deaminase domain may be unrelated to optimal catalytic competence.

Notably, one of the ADAR1 residues that interacts with C6-phosphate, N907, is associated with AGS. We show that the N907S mutation does not impair ADAR1 folding but markedly reduces RNA editing activity, consistent with AGS-associated loss-of-function variants (*67*).

Substitution of asparagine with serine introduces increased local dynamics within the hydrogen-bond interaction network, which is likely to propagate and disrupt precise Zn²⁺-ion positioning in the catalytic center. Ultimately, reduced RNA editing resulting from this mutation may facilitate aberrant MDA5 activation, thereby contributing to the pathogenesis of AGS.

Pro-IP_6_ treatment also enhanced RNA editing in wild-type cells, suggesting potential for modulating ADAR1 activity therapeutically. In the context of AGS, increased RNA editing could reduce aberrant MDA5 activation and thereby mitigate disease-associated inflammation.

In conclusion, several factors have been reported to regulate RNA editing, including ADAR dimerization (*26*), SUMOylation (*23*), and acidic pH (*25*). In this study, we demonstrate that IP_6_ is not only required for ADAR1 folding and RNA editing activity, but that its cellular levels also dictate the extent of editing, by increasing the amount of ADAR1. This finding suggests that IP_6_ levels represents an additional regulatory layer controlling RNA editing by ADAR proteins and raises important questions regarding the physiological contexts in which IP_6_ levels might fluctuate. Moreover, Pro-IP_6_ can enhance RNA editing, indicating that this molecule could have therapeutic potential in disorders such as AGS or for harnessing endogenous ADAR1 to correct pathogenic RNA mutations. Finally, our results indicate that the N907S mutation introduces an undesired dynamic into the hydrogen-bond interaction network linking IP_6_ to the ADAR1 catalytic center, thereby reducing RNA editing and providing mechanistic insight into how this disease-associated variant impairs ADAR1 function.

## MATERIALS AND METHODS

### Cell lines

HeLa WT and IPPK KO cells were maintained at 37 °C and 5% CO_2_ in RPMI 1640 medium supplemented with 10% FCS, 2 mmol/L L-glutamine, 100 U/mL penicillin G and 0.1% 2-mercaptoethanol.

Human embryonic kidney cells (HEK-293) WT and MINPP1 KO were obtained from Adolfo Saiardi (*55*), and cultivated at 37 °C and 5% CO_2_ in Dulbecco’s modified Eagle’s medium (DMEM) supplemented with 10% FCS, 2 mmol/L L-glutamine, 100 U/mL penicillin G and 0.1% 2-mercaptoethanol.

### Primary cells

Human peripheral blood mononuclear cells (hPBMCs) were isolated by density centrifugation using Lymphocyte separation media (anprotec) from buffy coats donated by anonymous healthy volunteers (the anti-coagulant used to obtain the buffy coats was sodium citrate). Informed consent was obtained from all blood donors. The local ethics committees of Justus-Liebig-University Gießen and Philipps-University Marburg approved the use of human blood samples (Az.: 24-67-BO).

### Generation of HeLa IPPK KO cells

The IPPK exon 2 targeting guide RNA (5′-AACAGCGCTGCGTCGTGCTG-3′) (*68*) was cloned into the vector pSpCas9(BB)-2A-Puro (PX459) (Addgene Plasmid # 48139), which was then transfected into HeLa cells using Lipofectamin 2000 (Cat. # 11668019, Thermo Fisher Scientific) followed by selection with puromycin at 2 μg/mL. IPPK deficient clones were obtained by limiting dilution and gene knockout was confirmed by western blotting and sequencing of a PCR-generated genomic fragment containing the targeting site using the primers 5’-GAAATGTGTGCCACTGTGTTTA-3′ and 5’-AGAAAGTGGTGTGTCCATCAT-3′ (*68*). The amplified fragment of IPPK knockout clone 1 contains a 10bp deletion and an unrelated sequence whereas clone 4 is characterized by a 2 and 10bp deletion (Fig. S13).

### RNA editing of specific targets in cells

RNA isolation, DNase digestion, and reverse transcription were performed according to the manufacturer’s protocol using TRIzol reagent (Cat. # 15596026, invitrogen), DNase I RNase-free (Cat. # 04716728001, Roche) and LunaScript RT Master Mix Kit (Cat. # E3025L, New England Biolabs), respectively. Subsequently a PCR was performed using specific primers for a fragment containing the resoective editing sites from *BLCAP*, *NEIL1*, *EEF2K* and *SON* (Table S4). PCR product was purified with GeneJET PCR Purification Kit (Cat. # K0701, Thermo Fisher Scientific) according to the manufacturer’s protocol and Sanger sequenced using standard forward or reverse primer of the specific primers for the editing sites (Seqlab, Microsynth). RNA editing quantification analysis was based on the area of adenosine and guanosine.

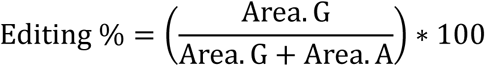

### Extraction of IPs and analysis

IPs were extracted according to Winson and Saiardi (*69*). Briefly, cell pellets were resuspended using 1 mol/L perchloric acid and incubated on ice for 10 min. Then, the samples were centrifuged (18,000 g for 5 min) and the supernatant was incubated with titanium beads (Cat. # 5020-75000, GL Sciences Inc.) (4 mg per sample) for 15 min at 4 °C with constant rotation. Subsequently, the beads were washed with 1 mol/L perchloric acid two times and resuspended with ammonium hydroxide 2.8% and incubated for 5 min at 4 °C with constant rotation. The supernatant of this extraction was evaporated using a centrifugal evaporator. These extracts were analyzed by Capillary Electrophoresis-Mass Spectrometry (CE-MS) or by a 36% Polyacrylamide Gel Electrophoresis (PAGE) gel with running TBE buffer and stained with a filtered staining solution of toluidine blue (20% methanol, 2% glycerol, 0.05% Toluidine blue) for 30 min. After destaining with washing solution (20% methanol, 2% glycerol), pictures were taken using a white light transilluminator.

For CE–MS analysis, IPs were extracted from 4×10⁶ cells for cell lines and 20×10⁶ cells for hPBMCs. Final extracts were resuspended in 20 µL prior to analysis.

For size-exclusion filtration experiments, hPBMC pellets (20 × 10⁶ cells) were lysed in Tris buffer containing 0.1% SDS, supplemented with a protease inhibitor cocktail (Cat. #78410, Thermo Fisher Scientific) and 5 mM EDTA. Lysates were incubated for 30 min at 4 °C under constant rotation and subsequently centrifuged at 13,000 × g for 10 min at 4 °C. The clarified supernatants were subjected to centrifugal ultrafiltration using Amicon® Ultra centrifugal filters with a 3 kDa molecular weight cut-off (MWCO). Both the filtrate and the retentate fractions were then processed for IP extraction using perchloric acid and titanium dioxide beads as described above. As a control for total IP content, 20×10⁶ hPBMCs were extracted directly using the standard perchloric acid/titanium bead protocol without prior filtration.

### Capillary Electrophoresis-Mass Spectrometry (CE-MS)

CE-MS measurements were performed on a CE-QqQ system for the analysis of InsP_5_ and InsP_6_ with internal standard [^13^C_6_] 1,3,4,5,6-IP_5_, [^13^C_6_] IP_6_ (*70*). All experiments were performed with a bare fused silica capillary with a length of 100 cm and 50 mm internal diameter. A 40 mM ammonium acetate, titrated with ammonia solution to pH 9.2, was used as background electrolyte. Samples were injected by applying 100 mbar pressure for 10 s (20 nL). The setting of the CE-QqQ system were the same as described in the literature (*32*, *34*).

### Stimulation of hPBMCs

80× 10^6^ hPBMCs were seeded in 25 cm^2^ cell culture flask incubated with IFN-α (1 ng/mL), LPS (1 ng/mL) and CpG-ODN 2216 (1 µM) or medium for 24 h in 7 mL RPMI medium supplemented with 2 mmol/L L-glutamine, 100 U/mL penicillin, 100 μg/mL streptomycin, 1 x amino acids, 1mM sodium pyruvate (PAN Biotech), 2% human AB serum (Biochrom AG). After this period of time, protein (10×10^6^ cells), RNA (5×10^6^ cells) and IPs (60×10^6^ cells) were extracted and analyzed.

### Western blotting

For HeLa cells and primary cells, cell pellet were lysed by lysis buffer (PBS-0.1% SDS) supplemented with Protease inhibitor cocktail (Cat. # 78410, Thermo Fisher Scientific) and 5 mM EDTA. Lysates were incubated for 30 min with constant rotation at 4 °C and subsequently centrifuged for 10 min at 13.000 g at 4 °C. Protein was quantified by BCA protein assay (Cat. # 23225, ThermoScientific) and 40 µg of the samples were then incubated at 95 °C for 3 minutes in SDS protein sample loading buffer and loaded and separated by tris-glycine SDS-PAGE followed by semiwet immunoblotting and antibody detection. Antibodies used were anti-ADAR1 (Cat. # sc-271854, Santa Cruz Biotechnology), anti-ADAR2 (Cat. # 67764-1-Ig, proteintech), anti-IPPK (Cat. # 12603-1-AP, proteintech) and anti-β-ACTIN (Cat. # A2228, Sigma-Aldrich) according to manufacturer’s instructions. For HEK293 cells, 30 µg protein was analyzed using anti-MINPP1 (Cat. # SC-514214, Santa Cruz,) and anti-GAPDH (Cat. # MA5-15738, Thermo Fisher Scientific) antibodies.

### Cloning of the ADAR1 editing reporter plasmid

The CDS of firely luciferase (Fluc) was excised from a plasmid via SalI and Eco72I and inserted downstream of a *Renilla* luciferase (Rluc) in a pMAX vector using the NEBuilder Hifi DNA Assembly Master Mix (Cat. # E2621, New England Biolabs). Additionally, a stem loop was inserted in-frame between the two luciferases via annealed oligonucleotides compromising homology arms. The stem loop, which has previously been used for constructing a similar reporter (*71*), contains an ADAR1 editing site within an UAG stop codon. Upon A-to-I editing, this is read as UGG and codes for tryptophan resulting in the translation of an Rluc-Fluc fusion protein.

### RNA editing–dependent dual-luciferase reporter assay (transient transfection)

HeLa or HEK293 cells were seeded in 96-well plates (30 000 cells/100 µL) and stimulated with Pro-IP_6_ or medium. After 2h medium was remplaced and cells were transiently transfected with the RNA editing reporter plasmid using Lipofectamine® 2000 (Thermo Fisher Scientific) according to the manufacturer’s instructions. Transfected cells were incubated for 24 h, then lysed with 50 µL Reporter Lysis Buffer (Cat. #E397A, Promega) per well. 20 µL of lysate was transferred to a Nunc F96 MicroWell white plate (Thermo Fisher Scientific) and enzymatic activities of Renilla and firefly luciferases were measured using the respective subtracts buffers and a Orion II Microplate Luminometer.

### Generation of *In vitro* translation ADAR plasmids

Human ADAR1 or ADAR2 were *in vitro* translated using PURExpress® In Vitro Protein Synthesis Kit (Cat. # E6800S, New England Biolabs) and the plasmids pIVTRA_ADAR1 (GenBank accession no. ON505744.1) or pIVTRA_ADAR2 (GenBank accession no. ON364011.1) coding for an N-terminal Twin-Streptag fused to ADAR1 (aa 432-931, NP_001020278.1) or to ADAR2 (aa 2-701, NP_001103). Both coding sequences were codon-optimized for expression in E. coli, synthesized as Strings™ DNA Fragments (GeneArt™ projects, Thermo Fisher Scientific) with 5’ and 3’ prime restriction sites NdeI and BamHI and cloned into the PURExpress control vector DHFR_Control_Template (Sequence online under https://international.neb.com/tools-and-resources/interactive-tools/dna-sequences-and-maps-tool) replacing the original DHFR gene. ADAR1 mutant constructs were generated by PCR-based site-directed mutagenesis (N907S, E912A and E1008A) or with a PCR fusion strategy (G1007R) with the primers listed in Table S5, followed by DpnI digestion to remove the parental plasmid template. Mutation in the plasmid template was confirmed by Sanger sequence.

### *In vitro* translation ADARs

*In vitro* translation of ADAR1 (ADAR1-IVTra) and ADAR2 (ADAR2-IVTra) was performed using the PURExpress In Vitro Protein Synthesis Kit (Cat. # E6800L, New England Biolabs) according to the manufacturer’s protocol. Briefly, reactions were assembled as follows: Solution A, Solution B, RiboLock RNase Inhibitor (Cat. # EO0382, Thermo Fisher Scientific), ZnCl₂ (final concentration 100 µM), ADAR-IVTra plasmid DNA (100 ng), H₂O (no IPs), and either IP_6_ (Cat. # P8810, Sigma-Aldrich) (1 µM), 1,3,4,5,6-IP_5_ (Cat. # sc-221502A, Santa Cruz Biotechnology) (1 µM). The mixture was gently mixed and incubated for 4 h at 27 °C. ADAR proteins were purified from the reaction mixture using MagStrep “type3” XT beads (Cat. # 2-4090-002, IBA) following the manufacturer’s instructions. The purified proteins were analyzed by SDS–PAGE followed by Coomassie staining or directly used for the *in vitro* RNA editing assay.

In the case of the ADAR1-IVTra with different IPs, the follow IPs were used in a final concentration of 0.1 µM, IP_6_ (Sigma-Aldrich), 1,3,4,5,6-IP_5_ (Santa Cruz Biotechnology), 1,2,6-IP_3_ (Cayman Chemical), 1,4,5-IP_3_ (Sigma-Aldrich), 1,3,4,5-IP_4_ (Sigma-Aldrich), 1,2,5,6-IP_4_ (Cayman Chemical), 2,3,4,5,6-IP_5_ (SiChem), 1,2,4,5,6-IP_5_ (SiChem), 1,2,3,5,6-IP_5_ (SiChem), 1,2,3,4,5-IP_5_ (SiChem).

### *In vitro* transcription (IVT)

Following DNA templates were synthesized as Strings™ DNA Fragments (GeneArt™ projects, Thermo Fisher Scientific) or amplified from genomic human DNA to generate ADAR1 and ADAR2 target RNA NEIL1, GLI1 or HTR2C by *in vitro* transcription: Human NEIL1 DNA template consists of 162 nucleotides (chromosome 15: 75353653-75353814, GRCh38.p14 assembly): 5’-GAGGTCTGGGCCAGGTCTAACCAGGCTCCCCACTCCTCCCAACCTGAGCCTGCCCTCTGATCTCTGCCTGTTCCTCTGTCCCACAGGGGGCAAAGGCTACGGGTCAGAGAGCGGGGAGGAGGACTTTGCTGCCTTTCGAGCCTGGCTGCGCTGCTATGGCAT-3′.

Using the primers NEIL1_F (5’-TAATACGACTCACTATAGGGAGGTCTGGGCCAGGTCTAAC-3′) and NEIL1_R (5’-ATGCCATAGCAGCGCAGC-3′) the fragment was amplified by PCR and thereby a T7 promoter introduced (adding two additional G to enhance *in vitro* transcription, sequence underlined).

The synthesized human GLI1 DNA template encompasses a T7 promotor with 3 G for optimal *in vitro* transcription (sequence underlined) and the GLI1 sequence (chromosome 12: 57470759-57470905, GRCh38.p13 assembly): 5’-TAATACGACTCACTATAGGGACAGAACTTTGATCCTTACCTCCCAACCTCTGTCTACTCACCACAGCCCCCCAGCATCACTGAGAATGCTGCCATGGATGCTAGAGGGCTACAGGAAGAGCCAGAAGTTGGGACCTCCATGGTGGGCAGTGGTCTGAACCCCTATAT-3′.

HTR2C DNA sequence was amplified from HEK293 genomic DNA using the primers HTR2C_F: 5’TAATACGACTCACTATAGGGCCCCGTCTGGATTTCTTTAGATG-3′ and HTR2C_R: 5’:CCGATCAAACGCAAATGTTACCAGTC-3′ and consists of 301 nt with a T7 promotor and 2 additional G for optimal *in vitro* transcription (sequence underlined) and the HTR2C genomic sequence (chromosome X:114848033-114848314, GRCh38.p13 assembly): 5’-TAATACGACTCACTATAGGGCCCCGTCTGGATTTCTTTAGATGTTTTATTTTCAACA GCGTCCATCATGCACCTCTGCGCTATATCGCTGGATCGGTATGTAGCAATACGTAATCCTATTGAGCATAGCCGTTTCAATTCGCGGACTAAGGCCATCATGAAGATTGCTATTGTTTGGGCAATTTCTATAGGTAAATAAAACTTTTTGGCCATAAGAATTGCAGCGGCTATGCTCAATACTTTCGGATTATGTACTGTGAACAACGTACAGACGTCGACTGGTAACATTTGCGTTTGATCGG-3′.

For *in vitro* transcription, 50 or 100 ng template DNA was transcribed with the T7-Scribe Standard RNA IVT Kit (Cellscript, Madison, USA) according to the manufacturer’s protocol. Briefly, the *in vitro* transcription reaction was incubated at 42 °C for 2 to 4 h, followed by template DNA digestion with DNase I for 30 min and then extracted with ROTI®Phenol/Chloroform/ Isoamyl alcohol (Cat. # A156.3, ROTH). RNA was desalted with BIO-RAD Micro-Spin Columns P-30 according to the manufacturer’s protocol. RNA size as well as integrity was analysed at 50 °C on a 10% (38:1) polyacrylamide gel containing 8 M Urea and 2x TAE buffer.

### Pro-IP_6_ treatment

Pro-IP_6_ was synthesized as previously described (*39*), or acquire from TCI (Cat. # P3062). HeLa WT and HeLa IPPK KO cells were seeded in 48-well plates (150 000 cells/500 µL) and incubated with Pro-IP_6_ at different concentrations, DMSO or medium for 24 h. After this period, RNA editing of *BLCAP* and *EEF2K* were evaluated. For WB analysis and IP_6_ analysis, 4×10^6^ cells were seeded in 25 cm^2^ cell culture flask in 7 mL with 10 µM Pro-IP_6_ or DMSO as a control and after 24 h, cells were harvested, 1×10^6^ cells were used for Western blotting analysis and 5 ×10^6^ cells were used for IP_6_ analysis.

### *In vitro* editing

IVT RNA (*GLI1*, *NEIL1* or *HTR2C*) was diluted to 2 ng/µL in reaction buffer containing 15 mM Tris-HCl (pH 7.6 at 25 °C), 60 mM KCl, 3 mM MgCl_2_, 1.5 mM DTT, 0.03 mM spermidine, heated to 95 °C for 3 min in a PCR cycler and cooled to 25 °C by reducing temperature every 2 min by 5 °C. Then, RNase Inhibitor RiboLock (1,6 U/µl) was added and 25 µL incubated with ADAR-IVTra (IP_6_, IP_5_ or no IPs) protein bound magnetic beads at 27 °C for 24 h.

For RT-PCR, 4 ng of IVT RNA were *in vitro* transcribed using the reverse primer of the respectively gene (Table S6) and QuantiTect Reverse Transcription Kit (Qiagen) or LunaScript RT Master Mix Kit (Cat. # E3025L, New England Biolabs) according to the manufacturer’s protocol. cDNA amplification was performed using the respectively primers from Table S6 and DreamTaq Green PCR Master Mix (2x) (Thermo Fisher Scientific). PCR product was purified with GeneJET PCR Purification Kit (Thermo Fisher Scientific) according to the manufacturer’s protocol and Sanger sequenced using standard T7 primer (5’-TAATACGACTCACTATAGGG-3′, Seqlab, Microsynth) or the reverse primer in the case of HTR2C (Table S6). Editing quantification analysis was based on the area of adenosine and guanosine.

### RNA processing and library preparation

HeLa WT and HeLa IPPK KO cells were seeded in 12-well plate (500 000 cells/1mL) and treated with Pro-IP_6_ 10 µM or DMSO as control for 24 h. After this period, total RNA was extracted using TRIzol reagent (Cat. #15596026, Invitrogen) according to the manufacturer’s instructions. Residual genomic DNA was removed with DNase I recombinant RNase-free (Cat. #04716728001, Roche), and RNA was subsequently purified using the RNA Clean & Concentrator-5 kit (Zymo Research).

All subsequent procedures, including RNA quality assessment, library preparation, and sequencing, were performed by GenomeScan BV (Leiden, the Netherlands). A total of 12 samples were received in good condition on August 25, 2025. RNA concentrations were determined using a fluorescence-based assay, and RNA integrity was evaluated with the QIAxcel Connect system.

Library preparation was carried out using the NEBNext Ultra II Directional RNA Library Prep Kit for Illumina (NEB #E7760S/L) following the manufacturer’s protocol. Briefly, mRNA was isolated from total RNA using oligo-dT magnetic beads, fragmented, and reverse transcribed into cDNA. After adapter ligation and PCR amplification, library quality and yield were verified using the QIAxcel Connect system.

Clustering and sequencing were performed on an Illumina NovaSeq X Plus platform according to the manufacturer’s guidelines, using a final loading concentration of approximately 160 pM DNA. Each sample was sequenced to a depth of at least 100 million reads. The dataset generated in this study is available in the NCBI Gene Expression Omnibus (GEO) under accession number GSE316528.

### Data analysis of NGS data

Sequencing reads from all lanes were merged at the FASTQ level prior to analysis. Quality filtering and trimming were performed with fastp (v0.23.4) with the following non-default parameters: -q 25 -u 20 -e 30 --dont_eval_duplication -G -A. Reads shorter than 97 bp were discarded (--length_required 97), and reads longer than 100 bp were trimmed from the 3′ end to this length (--max_len1 100). Transcript- and gene-level expression quantification was performed with Salmon (v1.10.2) using automatic library detection (-l A) and transcript-to-gene mapping (-g) based on a curated reference annotation. Reads were aligned to the human genome (hg38) with STAR (v2.7.3a) using parameters --alignSJoverhangMin 8 --alignIntronMax 1000000 --alignMatesGapMax 600000 --outFilterMultimapNmax 1 to ensure unique alignments.

### Differential expression and gene enrichment analysis

Differential expression analysis was performed on raw Salmon counts using DESeq2 (v1.48.1) in combination with tximport (v1.36.1) in R (v4.5.1). Genes with fewer than ten total counts across all samples were excluded. Log₂ fold changes were shrunk using the adaptive shrinkage (“ashr”) method to reduce noise from low-count genes. Significance was determined using false discovery rate (FDR)–adjusted p-values (Benjamini–Hochberg correction).

Genes with an adjusted *P* value < 0.05 and an absolute log₂ fold change > 1 were considered significantly differentially expressed and were used for downstream enrichment analyses. Gene symbols were converted to Entrez Gene IDs using the *org.Hs.eg.db* annotation package. Gene Ontology (GO) enrichment analysis for Biological Process (BP) terms and Kyoto Encyclopedia of Genes and Genomes (KEGG) pathway enrichment analysis were performed using the *clusterProfiler* package. Enrichment significance was assessed using a hypergeometric test, with *P* values adjusted for multiple testing using the Benjamini–Hochberg method. GO terms with an FDR (*q* value) < 0.05 and KEGG pathways with *P* < 0.05 were considered significantly enriched.

### Quantification A-to-I RNA editing levels

Global A-to-I RNA editing was quantified from STAR alignments using RNA Editing Indexer (v1.0) with the --genome hg38 flag to compute the *Alu* Editing Index (AEI). This index represents the mean editing level across all adenosines within *Alu* elements, weighted by their expression, and corresponds to the ratio of A-to-G mismatches to total aligned nucleotides in these regions — providing a robust global measure of A-to-I editing activity.

The Inverted *Alu* 3′UTR Editing Index was calculated similarly over inverted *Alu* clusters (-rb), defined as 3′UTR exons containing at least two *Alu* elements (each >200 bp) oriented on opposite strands. This metric captures editing activity within double-stranded RNA–forming 3′UTR structures, serving as a proxy for cytoplasmic A-to-I editing potential.

Site-specific RNA editing levels of the dataset of annotated editing sites described by Gabay et al. (*48*) were quantified with REDItools 1.0 in known-site mode with the following non-default parameters: -v 1 -n 0.001 -r 1 -T5-5 -c 1.

### Molecular dynamics simulations

MD simulations of deaminase domain (839-1223aa) of ADAR1-IP_5_/_6_ or ADAR1-N907S were performed based on the structure protein–ligand complex derived from PDB structure (PDB: 9B83). The protein was pre-processed using cpptraj (*72*) and standardized with pdb4amber. Ligand protonation and geometry optimization were performed using OpenBabel and RDKit with the MMFF force field. The complex was assembled and visually inspected prior to simulation setup. All atomistic MD simulations were carried out using OpenMM (*73*, *74*). The protein was parameterized with the ff19SB force field and the ligand with GAFF2, with AM1-BCC charges assigned via Antechamber. The system was solvated in a cubic TIP3P water box with a minimum solute–box distance of 10 Å, and NaCl was added to neutralize the system and to reach 0.15 M ionic strength. Periodic boundary conditions were applied in all dimensions.

Energy minimization was performed for 20,000 steps, followed by 0.5 ns NPT equilibration at 298 K and 1 bar using a Langevin thermostat and Monte Carlo barostat, respectively, with a 2 fs timestep. Positional restraints (700 kJ/mol) were applied to protein heavy atoms during equilibration. Production MD was performed for a total of 50 ns, executed as 10 consecutive strides of 5 ns each, using identical ensemble and integration settings. Trajectories were saved every 10 ps. Trajectory strides were concatenated using PyTraj (*72*), and water molecules were optionally removed prior to analysis. MM-GBSA binding free energies were computed using the igb = 5 model and 0.15 M salt concentration. Protein–ligand interaction energies, Cα RMSD, and Cα RMSF were calculated using PyTraj and MDAnalysis. Hydrogen bonds and interaction distances where visualized and analyzed using ChimeraX (*75*, *76*).

### Statistical analysis

Statistical significance was analysed using one-way ANOVA. When a significant effect was found, Dunnett’s or Tukey’s post-hoc test was used to contrast the effect of treatment with the control. In the case of two groups, a Student’s t test was used. Data analyses were performed using R software. For correlation analysis, Pearson correlation coefficient and matrix correlation based on Spearman correlation coefficient were performed using R software. A global P value lower than 0.05 was considered statistically significant (*P < 0.05, **P < 0.01, ***P < 0.001). Principal component analysis (PCA) was performed using R software.

## Acknowledgements

We thank G. Bein, Institute for Clinical Immunology and Transfusion Medicine, Justus-Liebig-University Giessen, for providing human buffy coats. We also thank Laila Awaragi for the help in the construction of editing reporter plasmid. FVS would like to thank Pedro LaRock for the very insightful conversations and opinions about RNA editing.

## Funding

This work was supported by Deutsche Forschungsgemeinschaft (DFG, German Research Foundation) Project-ID 369799452— TRR237 - A02 to SB and TZ, Project-ID 548714673 to FVS and CIBSS – EXC-2189 – Project ID 390939984 to HJ.

HJ and ML acknowledge funding from the Volkswagen Foundation (VW Momentum Grant 98604).

EYL was supported by the Israel Science Foundation grant 2637/23.

AS would like to acknowledge UKRI and the Medical Research Council for grant support (MR/T028904/1).

## Author contributions

Conceptualization: FVS, AK and SB. Project administration: FVS and SB Funding acquisition: FVS, HJ, AS, TZ, EYL and SB. Investigation: FVS, ML, RCF, FB, VF and SB. Formal analysis: FVS, ML, RCF, VF and SS. Visualization: FVS and SS.

Validation: FVS. Methodology: FVS, AK, HJ, AS, FB, EYL and SB. Writing—original draft: FVS. Writing—review and editing: FVS, ML, RCF, FB, VF, TM, MF, TZ, SS, AK, AS, EYL, HJ and SB. Supervision: FVS, AK, TZ, AS, EYL, HJ and SB. Resources: TM, MF, HJ and AS. Data curation and software: FVS, RCF and EYL.

## Competing interests

Authors declare that they have no competing interests.

## Data and materials availability

All data are available in the main text or the supplementary materials. The dataset generated in this study is available in the NCBI Gene Expression Omnibus (GEO) under accession number GSE316528.

## Notes

### Competing Interest Statement

The authors have declared no competing interest.

