## Supplementary Materials for "Inositol hexakisphosphate Functions as a Cofactor and Modulator of ADAR1 Activity"

### Inositol Hexakisphosphate as a Regulator of ADAR1-Mediated A-to-I RNA Editing

Francisco Venegas-Solis et al.

This PDF file includes:

Fig. S1 to S13

Table S1 to S6

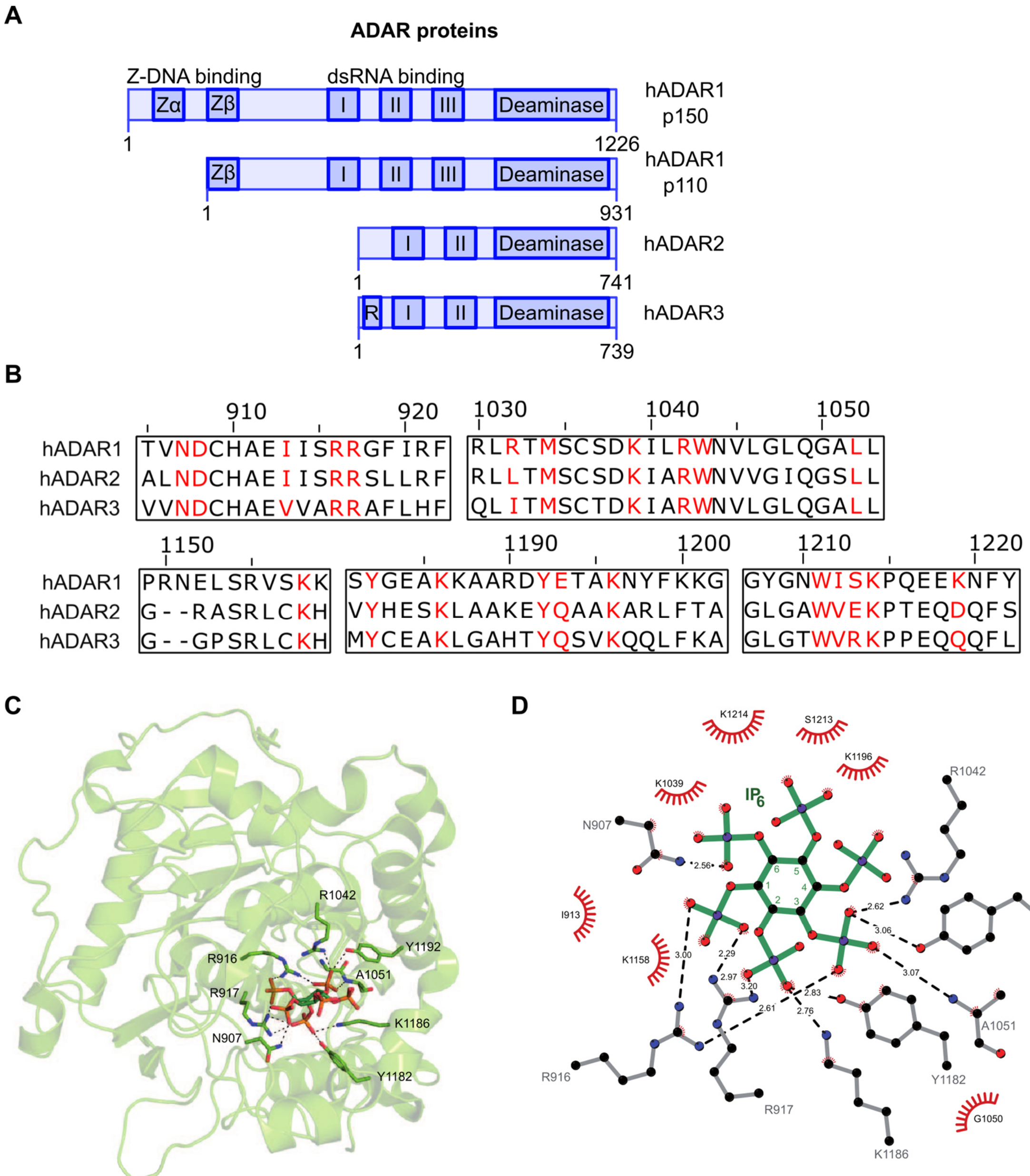


**Fig. S1. ADAR1-3 and IP$_6$ binding.** (**A**) Schematic representation of the domains of human ADAR1 (p150 and p110 isoforms), ADAR2 and ADAR3. Domains from left to right: Z-DNA binding domain (only ADAR1), double-stranded RNA binding (dsRNA binding), deaminase, R-rich domain (ADAR3 only). (**B**) Alignment of regions surrounding the residues that interact with IP$_6$ in ADAR1-3 (NP_001102.3, NP_001103.1 and NP_061172.1). IP$_6$ interacting residues are depicted in red. (**C**) Structure of ADAR1 in complex with IP$_6$ (PDB 9B83), with residues directly interacting with IP$_6$ (dark green) highlighted as stick representation. (**D**) Ligplot analysis of vicinity of the IP$_6$-binding site of ADAR1 (PDB 9B83).

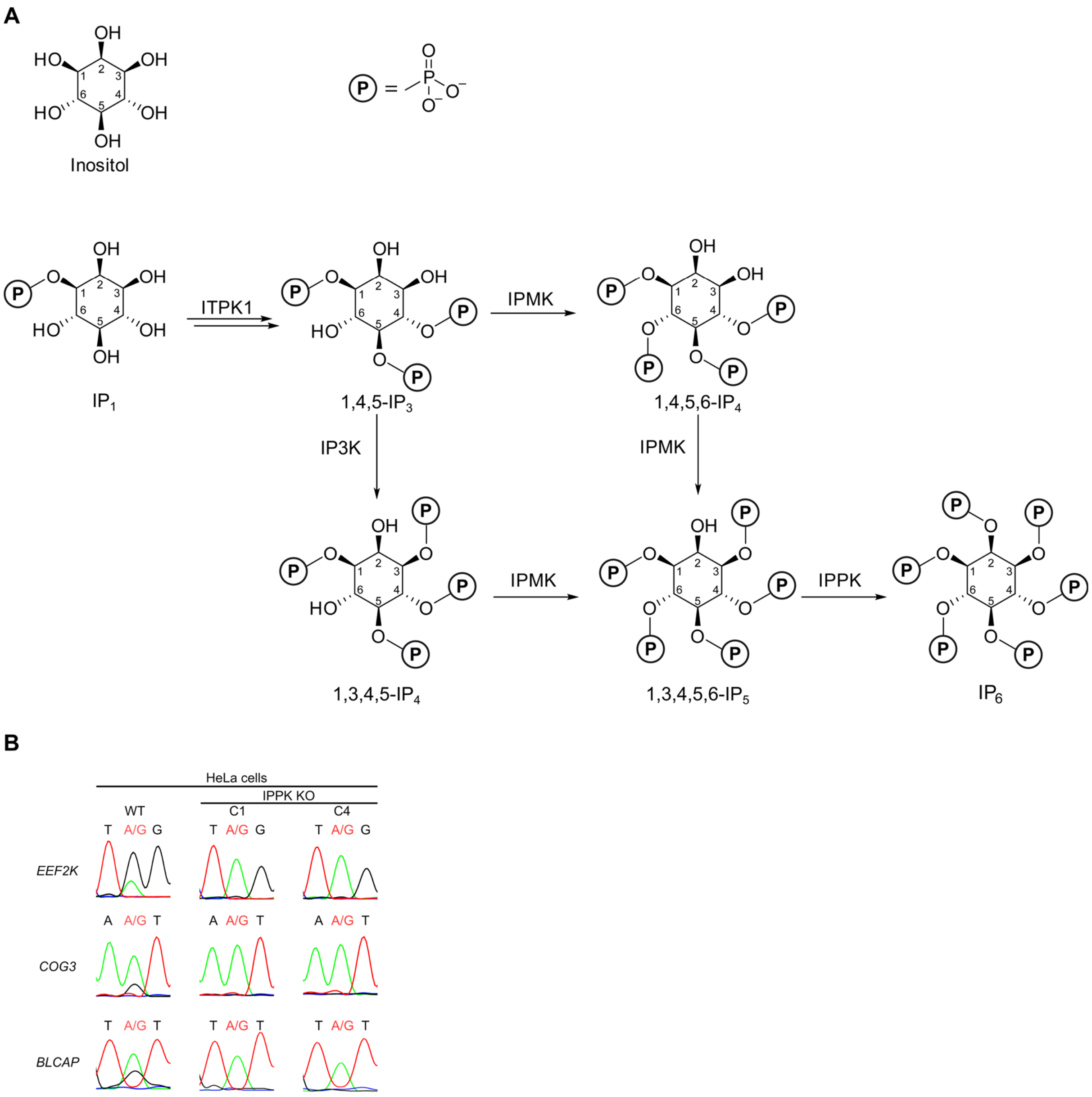


**Fig. S2. $IP_6$-byosynthesis.** (**A**) $IP_6$ biosynthesis pathway mediated by inositol-tetrakisphosphate 1-kinase (ITPK1), Inositol polyphosphate multikinase (IPMK), Inositol-trisphosphate 3-kinase A (IP3K) and Inositol-pentakisphosphate 2-kinase (IPPK). (**B**) Example of editing sequences of HeLa WT cells and HeLa KO IPPK cells clone 1 and 4 (C1 and C4).

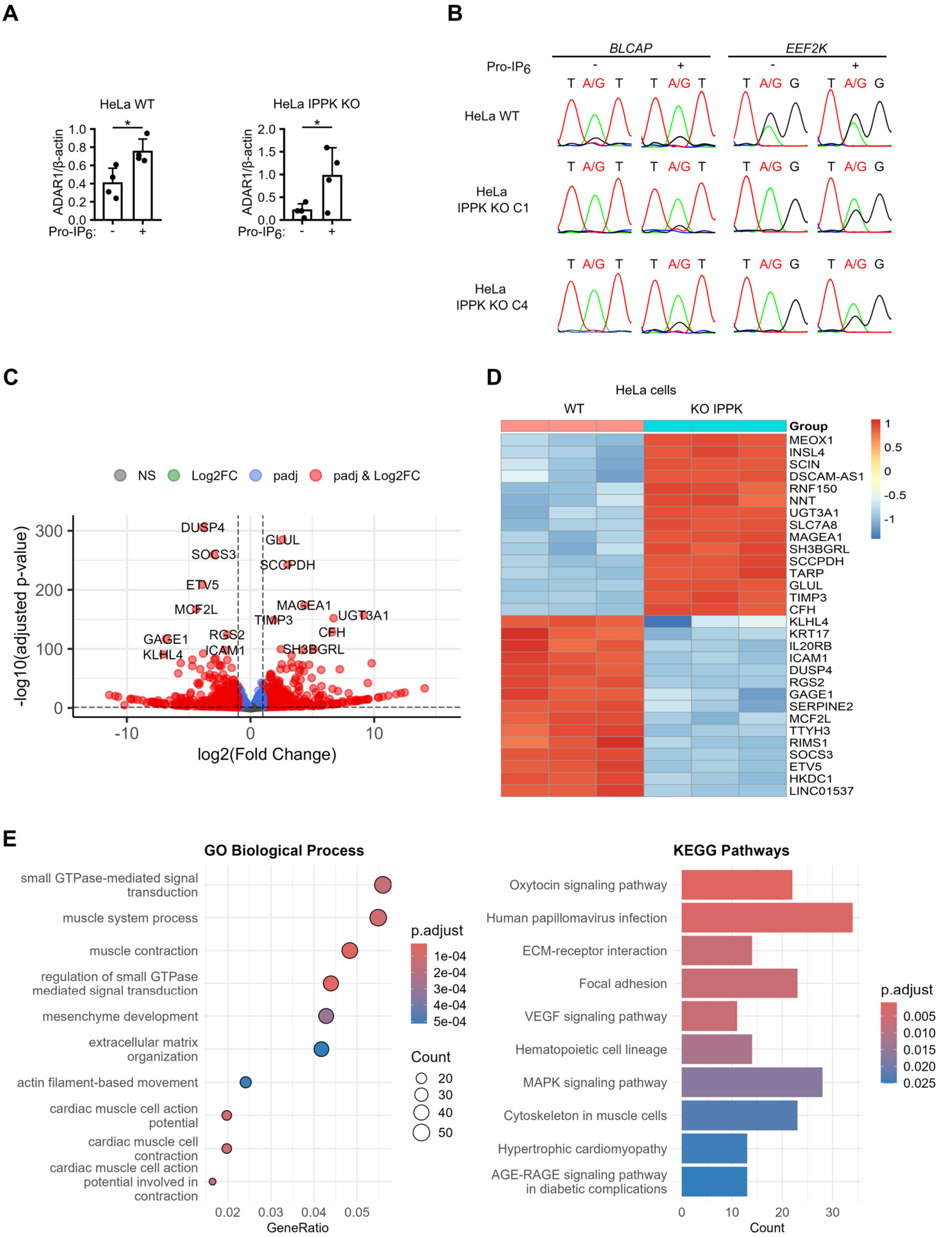


**Fig. S3. Transcriptomic profiling of HeLa WT and IPPK KO cells.** (**A**) Example of editing sequences of HeLa WT cells and HeLa KO IPPK cells (C1 and C4) untreated and treated with 10 µM Pro-IP$_6$. (**B**) Western blot quantification (n=4) of ADAR1 (p150+p110) and β-actin in HeLa WT cells and HeLa KO IPPK cells (C4) untreated and treated with 10 µM Pro-IP$_6$. (**C**) Volcano plot showing differentially expressed genes between HeLa WT and IPPK KO cells. (**D**) Heatmap of the top 30 differentially expressed genes. (**E**) GO enrichment and KEGG pathway enrichment analysis. Statistical significance was analyzed using Student’s t test, *$P < 0.05$, **$P < 0.01$, or ***$P < 0.001$.

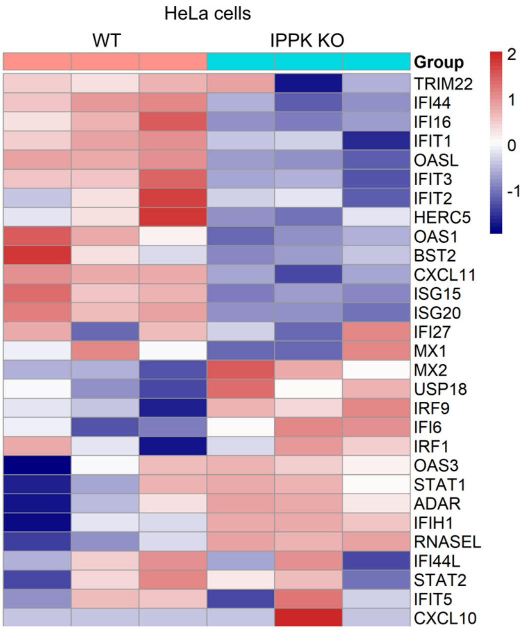


**Fig. S4. ISG expression in HeLa IPPK KO cells.** Heatmap of the top 20 differentially expressed ISG in HeLa WT and HeLa KO IPPK (C4) cells.

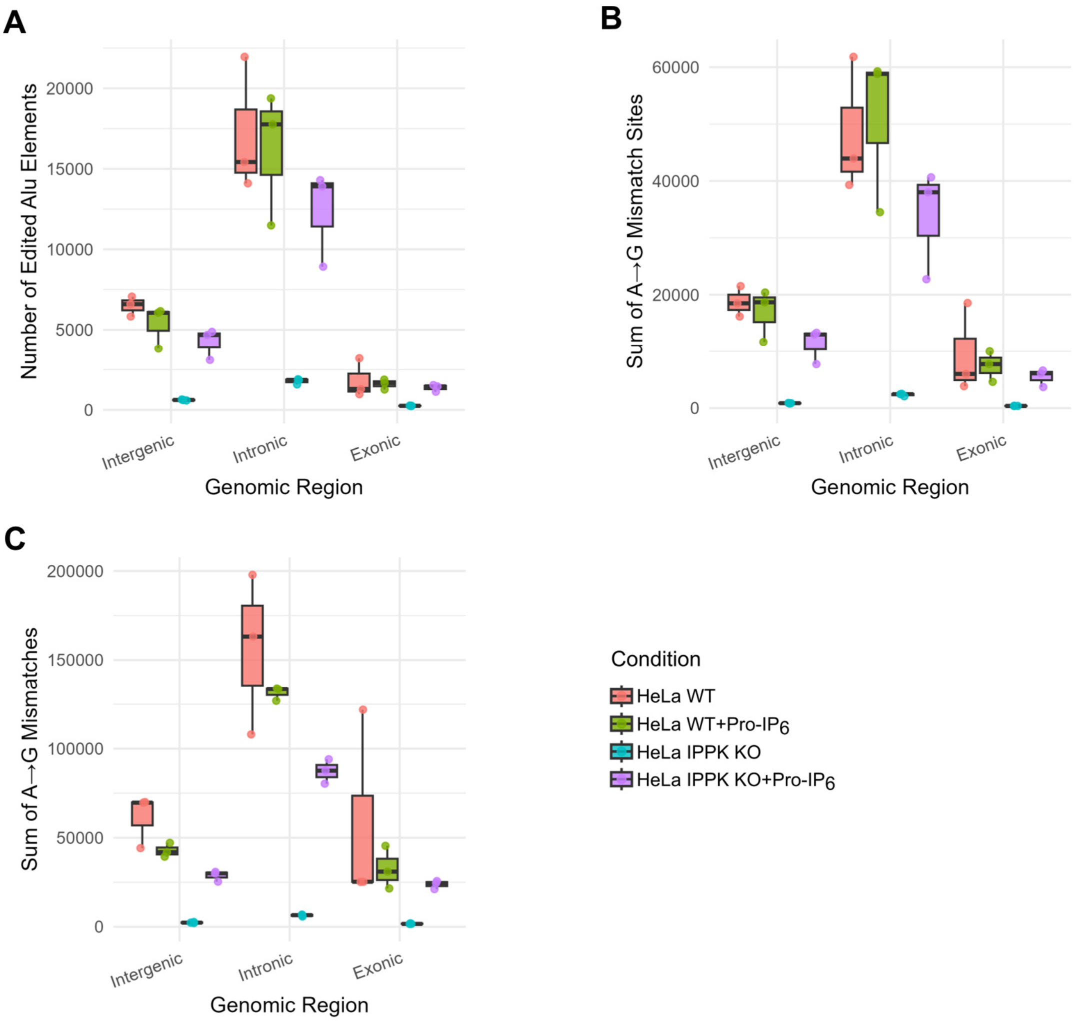


**Fig. S5. Analysis of A-to-I RNA editing across genomic regions.** Boxplots show the number of (**A**) edited *Alu* elements, (**B**) A→G mismatch sites, and (**C**) total A→G mismatches detected in intergenic, intronic, and exonic regions from wild-type HeLa cells (WT), WT HeLa cells treated with Pro-IP$_6$ (WT +Pro-IP$_6$), HeLa IPPK knockout (KO IPPK) cells, and HeLa IPPK KO cells treated with Pro-IP$_6$ (KO IPPK +Pro-IP$_6$). Each data point represents one experiment.

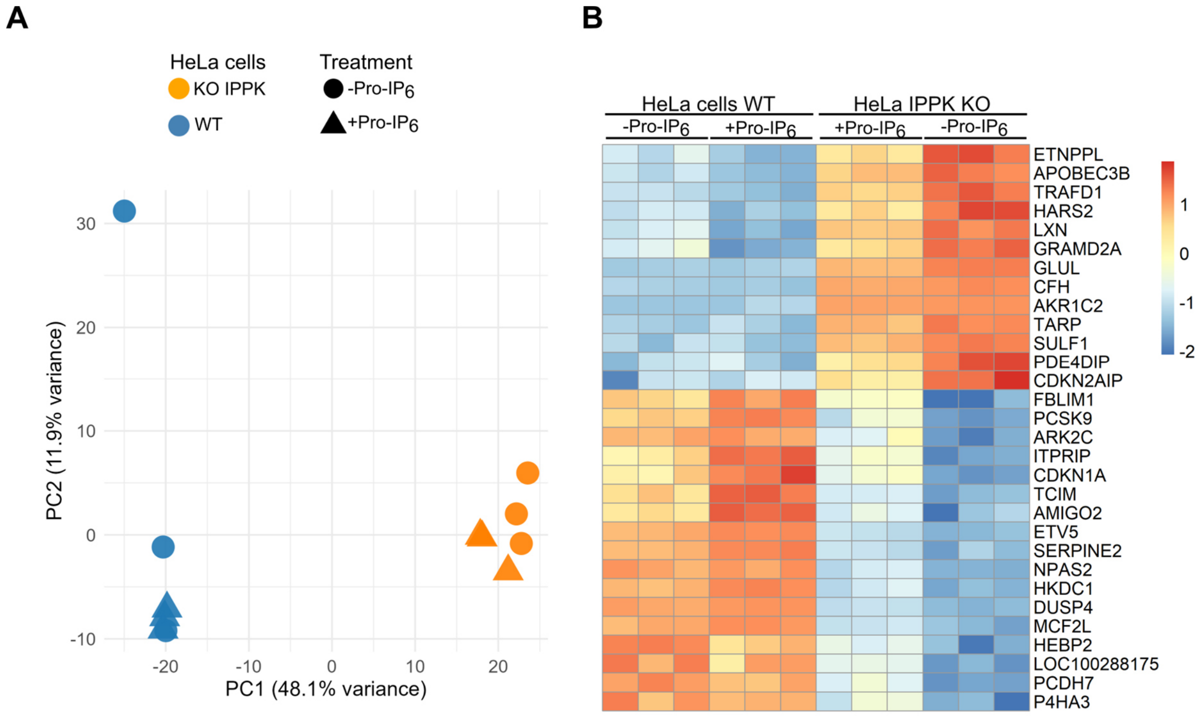


**Fig. S6. Pro-IP$_6$ effect on HeLa IPPK KO cells.** (**A**) PCA of the NGS data from HeLa WT and HeLa IPPK KO cells untreated or treated with Pro-IP$_6$. (**B**) Heatmap showing the expression of 30 genes downregulated or upregulated in HeLa IPPK KO cells relative to HeLa WT cells, which trend toward WT expression levels upon ProIP$_6$ treatment. Rows are scaled by z-score; columns correspond to biological replicates grouped by genotype and treatment. Annotation bars indicate genotype (WT = blue, KO = orange) and treatment (none = gray, Pro-IP$_6$ = green).

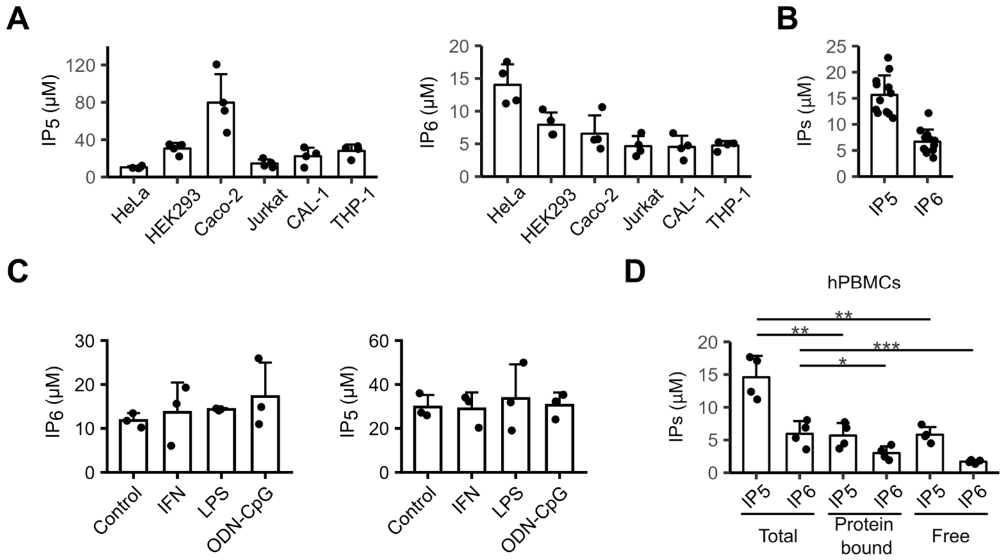


**Fig. S7. $IP_6$ and $IP_5$ protein bound or free**. (**A** and **B**) $IP_6$ and $IP_5$ (2-OH-$IP_5$) levels in different cell lines (n=4) or hPBMCs were analyzed by CE-MS. (**C**) Quantification of $IP_6$ and $IP_5$ in hPBMCs (n=3) (n=3) treated or not with IFN-α (1 ng/mL), LPS (1 ng/mL) and CpG-ODN 2216 (1 μM) for 20 h, and $IP_6$ levels were quantified by CE-MS. (**D**) Quantification of $IP_6$ and $IP_5$ from cell pellet, >3 kDa extract fraction (protein bound) and <3 kDa extract fraction (free) (n=4) were analyzed by CE-MS. Each data point represents one donor or one experiment. Statistical significance was analyzed using one-way ANOVA with Tukey's post hoc test, $^*P < 0.05$, $^{**}P < 0.01$, or $^{***}P < 0.001$ compared to total $IP_6/_5$.

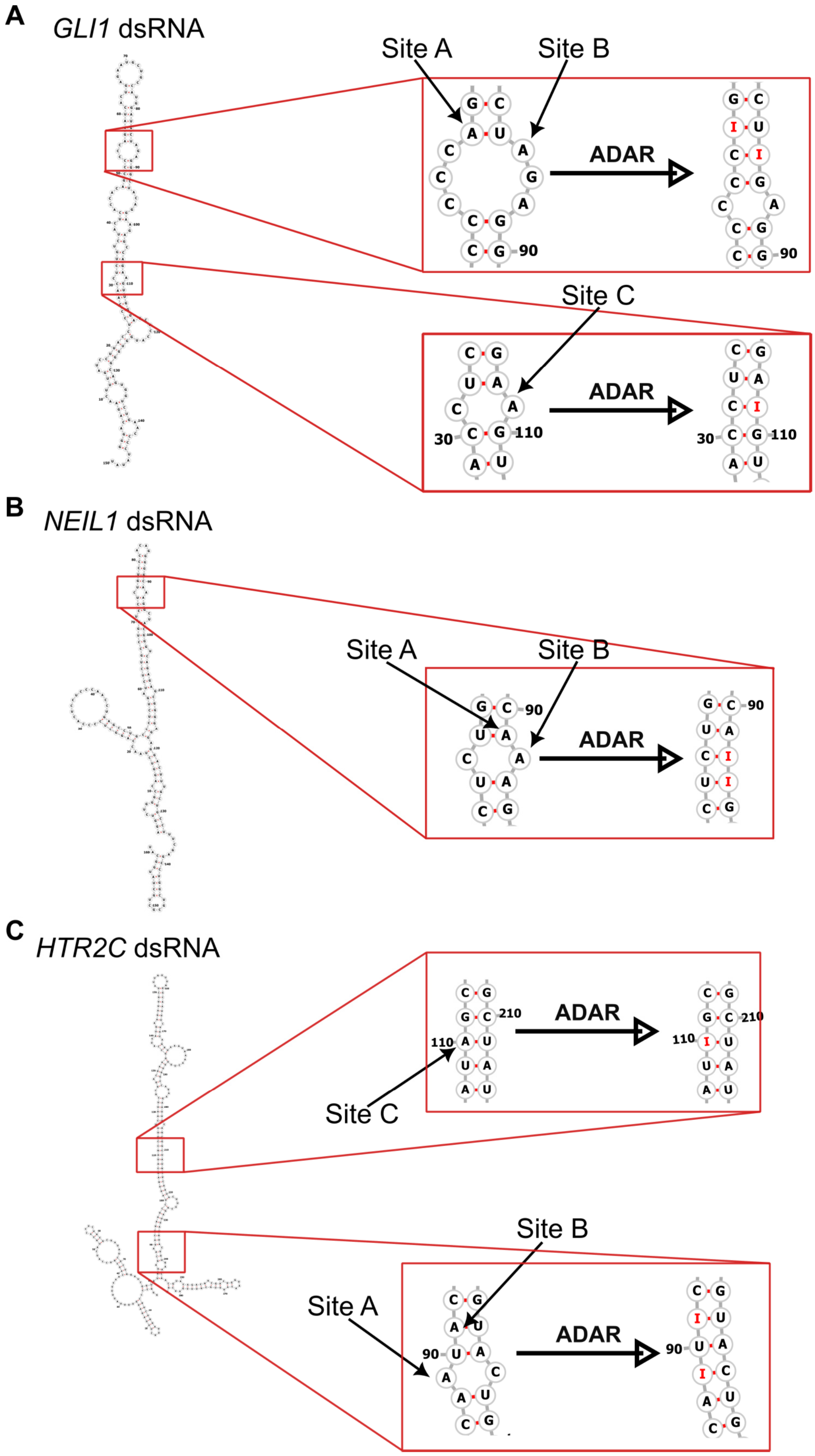


**Fig. S8. Secondary structures of *GLI1*, *NEIL1* and *HT2RC* dsRNA fragments.** Secondary structure prediction of *GLI1* (**A**), *NEIL1* (**B**) and *HT2RC* (**C**) dsRNA by RNAfold and visualization of editing sites as well as editing mediated changes in dsRNA structure.

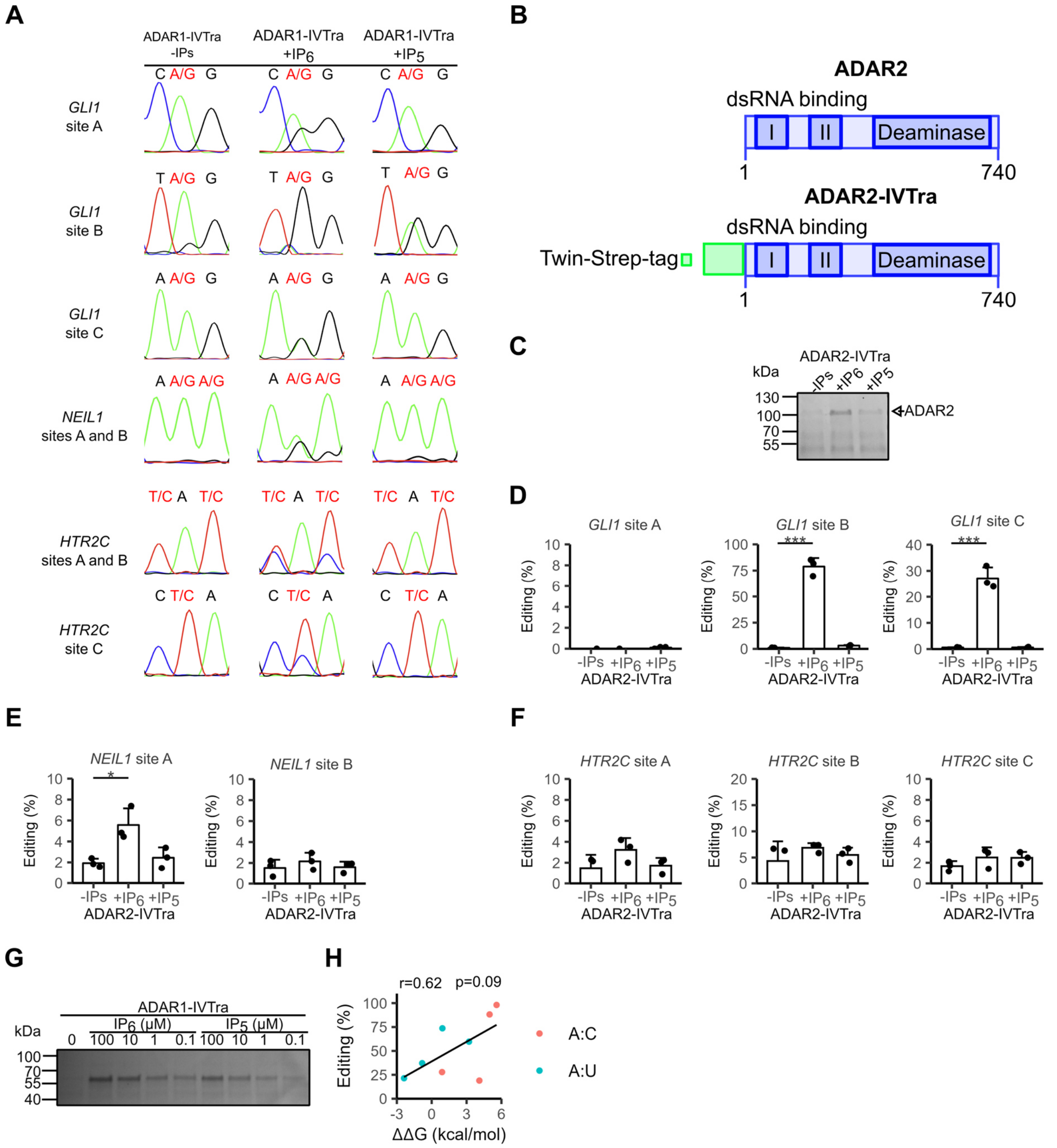


**Fig. S9. *In vitro* editing by *in vitro* ADAR2.** (**A**) Example of editing sequences of *in vitro* editing reactions using ADAR1-IVTra (no IPs, $IP_6$ and $IP_5$ as cofactors) and *in vitro* transcribed GLI1, NEIL1 and HTR2C dsRNAs. (**B**) Scheme of ADAR2 gene and ADA2 *in vitro* translated (IVTra) construct. (**C**) ADAR2-IVtra proteins used in the *in vitro* reactions of (**D-F**) were analyzed by SDS-PAGE and Coomassie staining. (**D-F**) *In vitro* editing reaction (n=3) of *in vitro* transcribed *GLI1*, *HTR2C* and *NEIL1* dsRNA using ADAR2-IVTra (no IPs, $IP_6$ and $IP_5$ as cofactors); editing levels were analyzed by cDNA synthesis follow up by PCR amplification and sanger sequencing. (**G**) Example of a gel used for the quantification of ADAR1-IVTra using different concentration of $IP_6$ and $IP_5$. (**H**) Correlation between % editing and ΔΔG that cause the changes of the editing in the secondary structure; ΔΔG was calculated using RNAfold before and after editing of the specific sites. Pink dots represent the mismatch (A:C) in the editing site and turquoise dots represent the prefect match (A:U). Each data point represents one experiment. For correlation analysis, Spearman correlation coefficients were performed, and a $P < 0.05$ was considered significant. Statistical significance was analyzed using one-way ANOVA with Tukey's post hoc test, $^{*}P < 0.05$, $^{**}P < 0.01$, or $^{***}P < 0.001$ compared to ADAR1-IVTra-no IPs.

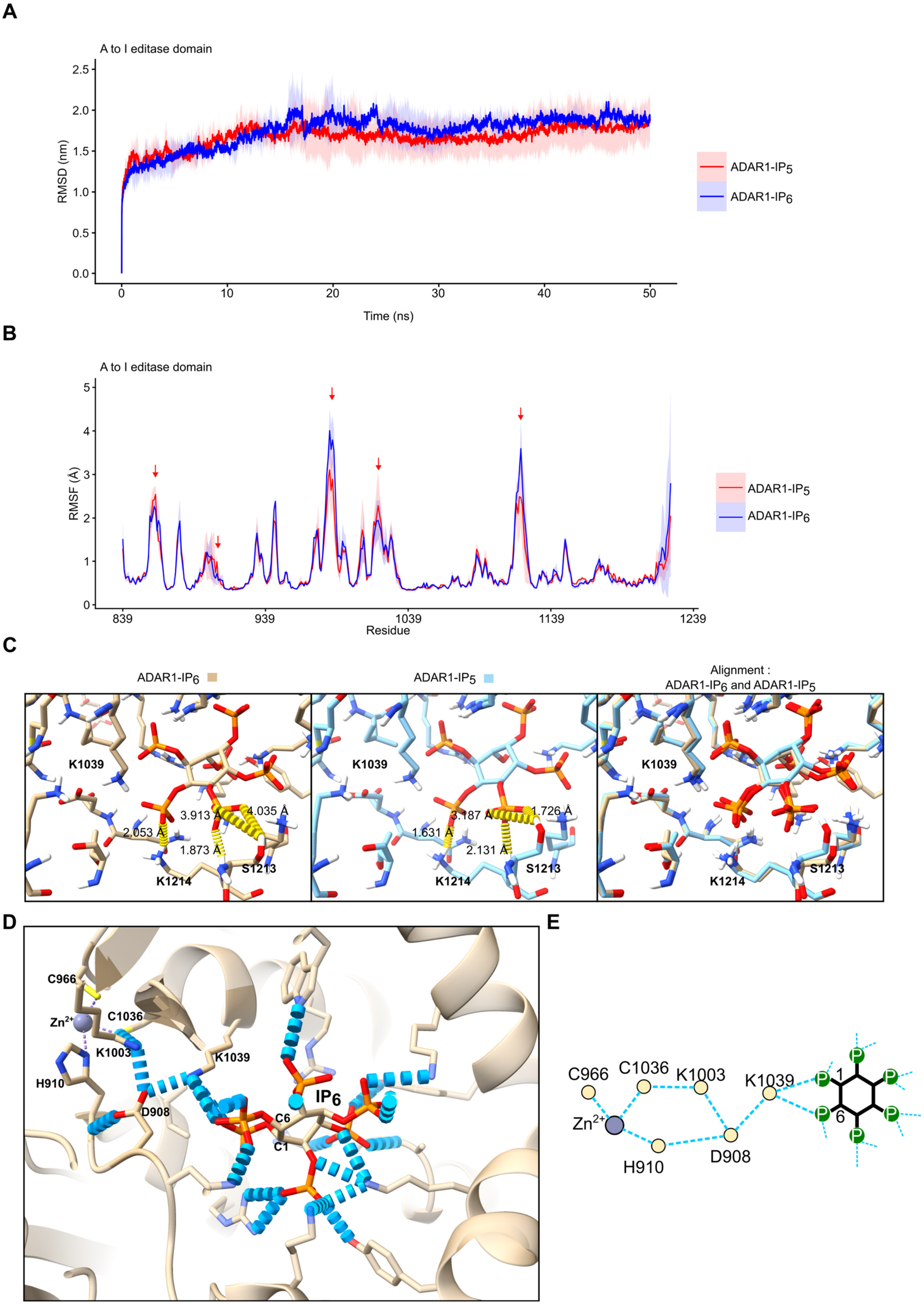


**Fig. S10. Molecular dynamics (MD) analysis of ADAR1–IP$_6$ and ADAR1–IP$_5$.** (**A**) Root-mean-square deviation (RMSD) and (**B**) root-mean-square fluctuation (RMSF) analyses of individual residues of the ADAR1 deaminase domain in complex with IP$_6$ or IP$_5$. Curves represent the mean of three independent simulations (n = 3); shaded areas indicate the standard deviation, and red arrows indicate visually apparent differences between the complexes. (**C**) Representative MD snapshots of ADAR1–$IP_6$, ADAR1–$IP_5$, and their alignment, showing the distances between S1213–C5-phosphate and K1214–C5/C6-phosphates. (**D**) Hydrogen-bond interaction network linking $IP_6$ to the ADAR1 catalytic center. (**E**) Simplified and schematic version of the hydrogen-bond interaction network linking $IP_6$ to the ADAR1 catalytic center.

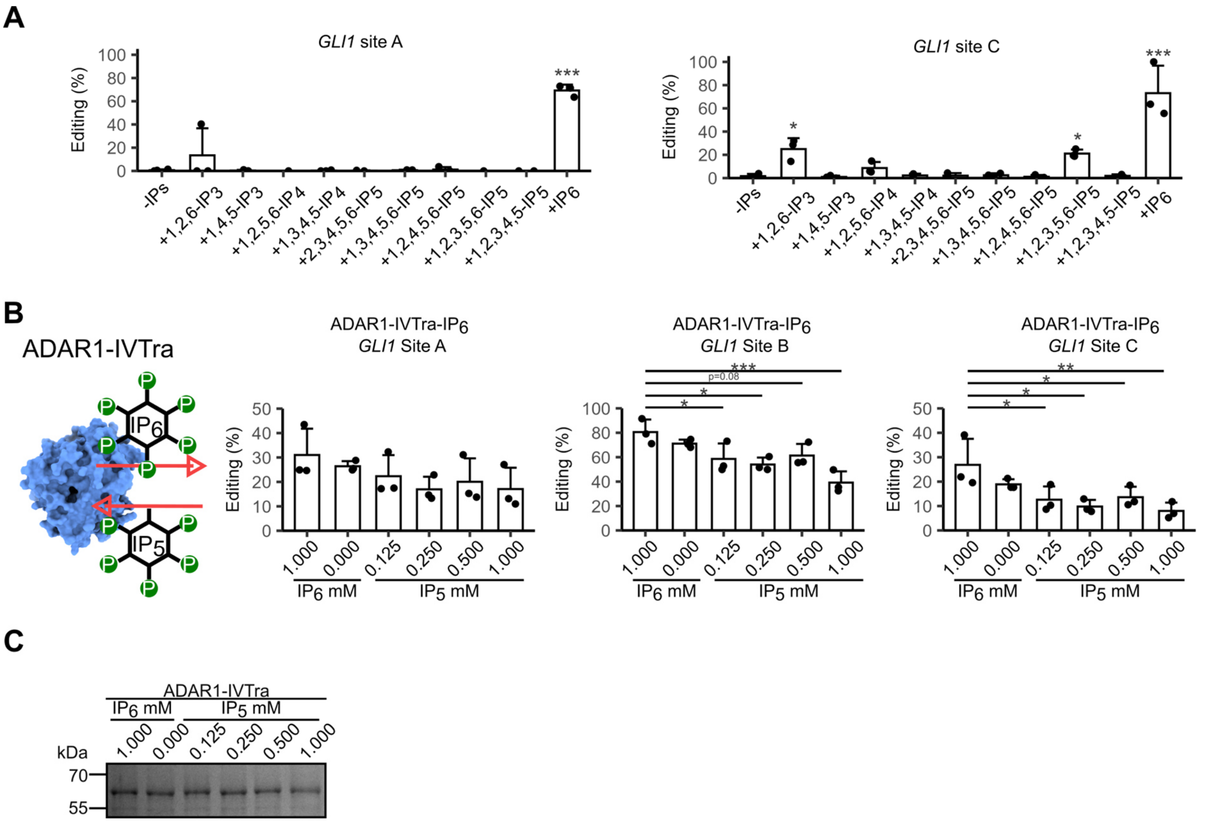


**Fig.S11. Interchange between IP$_6$ and IP$_5$ in ADAR1.** (**A**) Schematic representation of the interchange between IP$_6$ and IP$_5$ in ADAR1 and editing analysis of *in vitro* with IP$_6$ generated ADAR IVTra on *GLI1* dsRNA (n=3) incubating the editing reaction with different IP$_5$ concentrations or, IP$_6$/no IPs as controls. (**B**) ADAR1-IVtra proteins used in the *in vitro* reactions of (A) were analyzed by SDS-PAGE and Coomassie staining. (**C**) *In vitro* editing reaction (n=3) of *GLI1* dsRNA site A and C using ADAR1-IVTra with different IPs as cofactors. Editing levels were analyzed by cDNA synthesis followed by target PCR amplification and sanger sequencing. Each data point represents one experiment. Statistical significance was analyzed using one-way ANOVA with Dunnett's post hoc test (G), $^*P < 0.05$, $^{**}P < 0.01$, or $^{***}P < 0.001$ compared to ADAR1-IVTra-IP$_6$ 1 mM (A) or ADAR1-IVTra-no IPs (C).

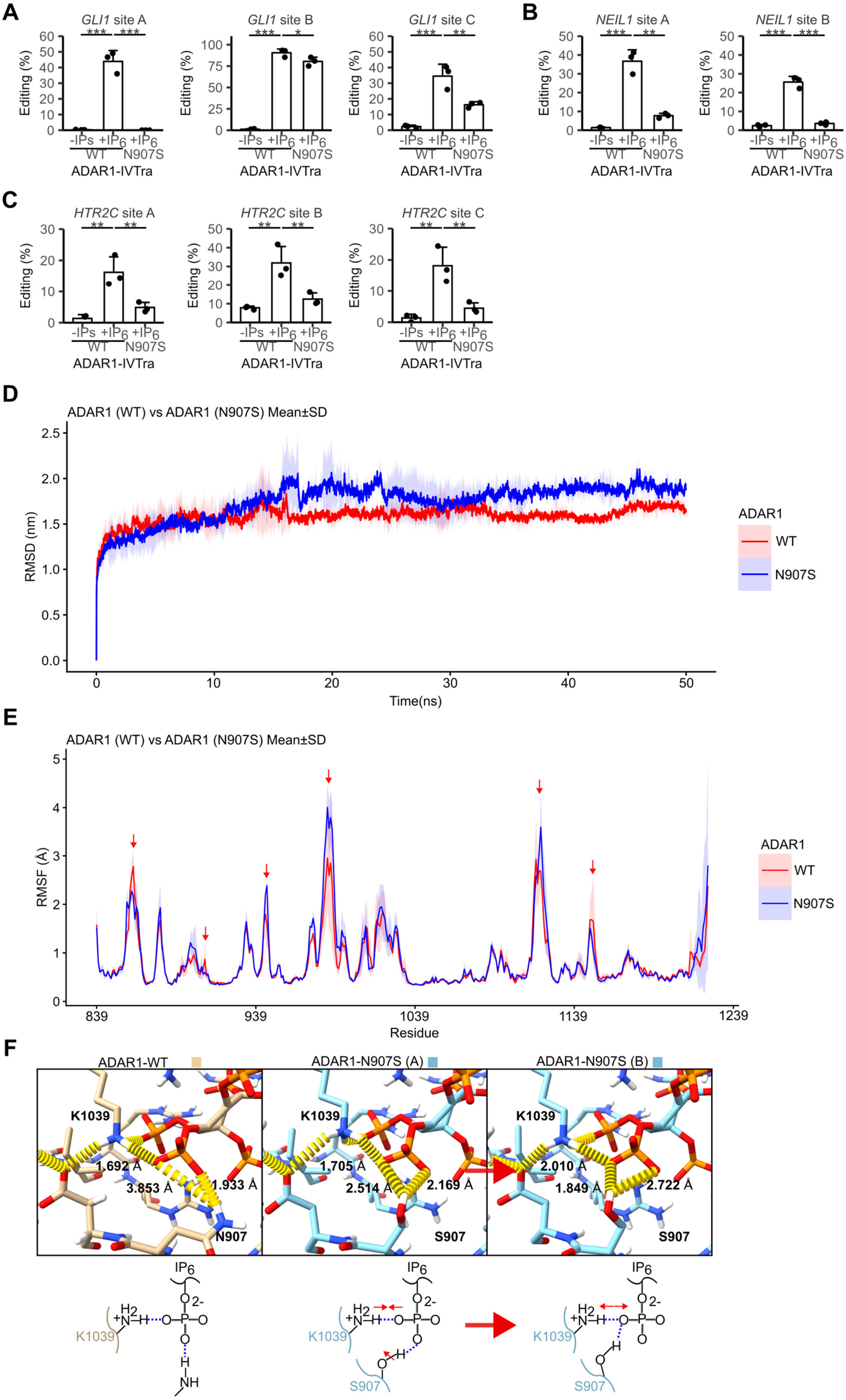
A
GLI1 site A
GLI1 site B
GLI1 site C
B
NEIL1 site A
NEIL1 site B
Editing (%)
-IPs +IP6 +IP6
WT N907S
ADAR1-IVTra
C
HTR2C site A
HTR2C site B
HTR2C site C
D
ADAR1 (WT) vs ADAR1 (N907S) Mean±SD
RMSD (nm)
Time(ns)
ADAR1
WT
N907S
E
ADAR1 (WT) vs ADAR1 (N907S) Mean±SD
RMSF (Å)
Residue
F
ADAR1-WT
ADAR1-N907S (A)
ADAR1-N907S (B)
K1039
N907
S907
1.692 Å
3.853 Å
1.933 Å
1.705 Å
2.514 Å
2.169 Å
2.010 Å
1.849 Å
2.722 Å
IP6

**Fig. S12. Editing profile of ADAR1 N907S mutant and MD analysis of ADAR1-WT and ADAR1-N907S.** *In vitro* editing reactions (n = 3) of *GLI1* (**A**), NEIL1 (**B**), and *HTR2C* (C) dsRNA substrates using ADAR1-IVTra (no IPs or $IP_6$ as cofactors) and ADAR1-IVTra N907S ($IP_6$ as cofactor). Editing levels were analyzed by cDNA synthesis followed by target PCR amplification and Sanger sequencing. (**D**) Root-mean-square deviation (RMSD) and (**E**) root-mean-square fluctuation (RMSF) analyses of individual residues of the ADAR1 deaminase domain for ADAR1-WT and ADAR1-N907S. Curves represent the mean of three independent simulations (n = 3); shaded areas indicate the standard deviation, and red arrows indicate visually apparent differences between the proteins. (**F**) Representative MD snapshots of ADAR1-WT and ADAR1-N907S showing the distances between N907/S907–C6-phosphate and K1039–C6-phosphate under the specific conformations observed. For panels (A–C), comparisons were made relative to ADAR1-IVTra-no IPs. *P < 0.05, **P < 0.01, or ***P < 0.001.

**A**

```
301 TTATTTTCCAACAGCGCTGCGTCGTGCTGCGGTTTCTGAAGTTTCCT
           ||||||||||||||||          ||||||||||||||||
1 -------TTCCAACAGCGCTGCG----------GTTTCTGAAGTTTCCT 83
```

**B**

```
301 TTTTATTTTCCAACAGCGCTGCGTCGTGCTGCGGTTTCTGAAGTTTCCTCCAAATAGGAAGAAGGTAAGATGGGTGGGTTTGCACCCAGACTTACAACAA 400
    TTTTATTTTCCAACAGCGCTGCGTC----------TTCTGAAGTTTCCTCC-------------------------------------------------  57
    TTTTATTTTCCAACAGCGCTGCG--GTGCTGCGGTTTCTGAAG
```

**Fig. S12. Detection of the alterations due to site-specific Cas9 activity.** (**A**) Sequence alignment of genomic fragments from WT IPPK and a sequenced PCR amplified fragment of IPPK KO clone 1 (C1) contains a 10bp deletion. (B) Sequence alignment of genomic fragments from WT IPPK and a sequenced PCR amplified fragment of IPPK KO clone 4 contains a 2 and 10bp deletion.

**Table S1. Location of gene specific editing sites**

| Gene | Chr | Site | Position | strand | Location | Region | Genome Reference |
|---|---|---|---|---|---|---|---|
| *BLCAP* | 20 | - | 37519170 | - | NONREP | exonic | GRCh38.p14 |
| *NEIL1* | 15 | A | 75353745 | - | NONREP | exonic | GRCh38.p14 |
| *NEIL1* | 15 | B | 75353746 | - | NONREP | exonic | GRCh38.p14 |
| *EEF2K* | 16 | - | 22285539 | + | ALU | UTR3 | GRCh38.p14 |
| *SON* | 21 | - | 33551013 | - | NONREP | exonic | GRCh38.p14 |
| *COG3* | 13 | - | 45516236 | - | NONREP | exonic | GRCh38.p14 |
| *GLI1* | 12 | A | 57470811 | + | NONREP | exonic | GRCh38.p14 |
| *GLI1* | 12 | B | 57470841 | + | NONREP | exonic | GRCh38.p14 |
| *GLI1* | 12 | C | 57470864 | + | NONREP | exonic | GRCh38.p14 |
| *HTR2C* | X | A | 114848119 | + | NONREP | exonic | GRCh38.p14 |
| *HTR2C* | X | B | 114848121 | + | NONREP | exonic | GRCh38.p14 |
| *HTR2C* | X | C | 114848140 | + | NONREP | exonic | GRCh38.p14 |

**Table S2.** *Alu* editing index and inverted *Alu* in 3'UTR editing index of HeLa cells (WT) and HeLa IPPK KO (C4) cells untreated or treated with10 µM Pro-$IP_6$

| Genotype | Pro-$IP_6$ | n | A to G editing index | sd A to G editing index | A to G editing index (Inverted *Alu* ) | sd A to G editing index (Inverted *Alu* ) |
|---|---|---|---|---|---|---|
| WT | - | 3 | 1,134 | 0,147 | 3,056 | 0,047 |
| WT | + | 3 | 1,320 | 0,055 | 3,053 | 0,126 |
| IPPK KO | - | 3 | 0,053 | 0,002 | 0,040 | 0,001 |
| IPPK KO | + | 3 | 0,950 | 0,025 | 2,125 | 0,057 |

sd: standard deviation

**Table S3. RNA editing activity across genomic regions in response to $IP_6$ treatment**

| Genomic region | Genotype | Pro-$IP_6$ | mean n edited *Alu* elements | sd n edited *Alu* elements | mean sum num of A to G Mismatches Sites | sd sum num of A to G Mismatches Sites | mean sum num of A T G Mismatches | sd sum num of A T G Mismatches | n |
|---|---|---|---|---|---|---|---|---|---|
| Intergenic | WT | - | 6497 | 625 | 18737 | 2679 | 61276 | 14821 | 3 |
| Intergenic | WT | + | 5347 | 1324 | 16924 | 4613 | 42779 | 3961 | 3 |
| Intergenic | IPPK KO | - | 614 | 29 | 864 | 53 | 2345 | 302 | 3 |
| Intergenic | IPPK KO | + | 4218 | 969 | 11357 | 3141 | 28699 | 2937 | 3 |
| Intronic | WT | - | 17160 | 4206 | 48358 | 11884 | 156345 | 45320 | 3 |
| Intronic | WT | + | 16209 | 4169 | 50867 | 14163 | 131469 | 3965 | 3 |
| Intronic | IPPK KO | - | 1781 | 174 | 2353 | 233 | 6373 | 552 | 3 |
| Intronic | IPPK KO | + | 12378 | 2998 | 33796 | 9677 | 87338 | 6857 | 3 |
| Exonic | WT | - | 1827 | 1208 | 9479 | 7935 | 57441 | 55825 | 3 |
| Exonic | WT | + | 1605 | 312 | 7450 | 2699 | 32639 | 12053 | 3 |
| Exonic | IPPK KO | - | 253 | 9 | 388 | 5 | 1711 | 189 | 3 |
| Exonic | IPPK KO | + | 1376 | 219 | 5478 | 1557 | 23768 | 2352 | 3 |

sd: standard deviation

**Table S4. Primers used for RNA editing of specific targets in cells**

| Gene | Forward primer (5'->3') | Reverse primer (5'->3') | Product length |
|---|---|---|---|
| *BLCAP* | CTGACAGCCAGAGAGCACAG | ATTGTGCAAGGTTTCCGTTC | 283 |
| *EEF2K* | CCCACAACGTGACTGCAATG | GGCTGCTTTTGCATTTAGGGT | 522 |
| *SON* | TGTGACGACGACAGAGTTGG | GTAGTAGGCAGCTCCTGTGC | 561 |
| *NEIL1* | ATGTCACTCAGTGCCCAAGG | CCTGGAACCAGATGGTACGG | 164 |
| *COG3* | TCTGGTTCAACAGAATCCCTCA | AACTGCTCACAGGCCGATTT | 509 |
| AluSx *CTSS* | GGCTCCTTCTCCATAAAGCA | AAAGTAGGCTGGGCTCAGTG | 442 |

**Table S5. Primers used in the mutagenesis**

| Mutation | Forward primer (5'->3') | Reverse primer (5'->3') |
|---|---|---|
| N907S | CCTGAAAGGTGAAACCGTTAGTGATTGTCAT GCCGAAATTA | TAATTTCGGCATGACAATCACTAACGGTTTC ACCTTTCAGG |
| E912A | TGATTGTCATGCCGCAATTATCAGCCGTC | GACGGCTGATAATTGCGGCATGACAATCA |
| E1008A | GTTGAAAATGGTGCAGGCACCATTCCG | CGGAATGGTGCCTGCACCATTTTCAAC |
| G1007R | ACGTAGCCATGGTTTTGCAGCAG | ACGATTTTCAACTTTGGTACGCAG |
| G1007R | AAAGTTGAAAATCGTGAAGGCACCATTCCG GTTGAA | ATATATGGATCCTTACACCGGACAC |

**Table S6. Primers used in the *in vitro* editing**

| Gen | Forward primer (5'->3') | Reverse primer (5'->3') |
|---|---|---|
| *GLI1* | TAATACGACTCACTATAGGGACAGAACTTTGATCCTTACCTC | ATATAGGGGTTCAGACCACTGCCCAC |
| *NEIL1* | TAATACGACTCACTATAGGGAGGTCTGGGCCAGGTCTAAC | ATGCCATAGCAGCGCAGC |
| *HTR2C* | TAATACGACTCACTATAGGGCCCCGTCTGGATTTCTTTAGATG | CCGATCAAACGCAAATGTTACCAGTC |
| T7 | TAATACGACTCACTATAGGG | |
